# Targeted, High-resolution RNA Sequencing of Non-coding Genomic Regions Associated with Neuropsychiatric Functions

**DOI:** 10.1101/539882

**Authors:** Simon A. Hardwick, Samuel D. Bassett, Dominik Kaczorowski, James Blackburn, Kirston Barton, Nenad Bartonicek, Shaun L. Carswell, Hagen U. Tilgner, Clement Loy, Glenda Halliday, Tim R. Mercer, Martin A. Smith, John S. Mattick

**Author notes:** These authors contributed equally (Simon A. Hardwick, Samuel D. Bassett).

## Abstract

The human brain is one of the last frontiers of biomedical research. Genome-wide association studies (GWAS) have succeeded in identifying thousands of haplotype blocks associated with a range of neuropsychiatric traits, including disorders such as schizophrenia, Alzheimer’s and Parkinson’s disease. However, the majority of single nucleotide polymorphisms (SNPs) that mark these haplotype blocks fall within non-coding regions of the genome, hindering their functional validation. While some of these GWAS loci may contain *cis-*acting regulatory DNA elements such as enhancers, we hypothesized that many are also transcribed into non-coding RNAs that are missing from publicly available transcriptome annotations. Here, we use targeted RNA capture (‘RNA CaptureSeq’) in combination with nanopore long-read cDNA sequencing to transcriptionally profile 1,023 haplotype blocks across the genome containing non-coding GWAS SNPs associated with neuropsychiatric traits, using post-mortem human brain tissue from three neurologically healthy donors. We find that the majority (62%) of targeted haplotype blocks, including 13% of intergenic blocks, are transcribed into novel, multi-exonic RNAs, most of which are not yet recorded in GENCODE annotations. We validated our findings with short-read RNA-seq, providing orthogonal confirmation of novel splice junctions and enabling a quantitative assessment of the long-read assemblies. Many novel transcripts are supported by independent evidence of transcription including cap analysis of gene expression (CAGE) data and epigenetic marks, and some show signs of potential functional roles. We present these transcriptomes as a preliminary atlas of non-coding transcription in human brain that can be used to connect neurological phenotypes with gene expression.

## INTRODUCTION

Over the past decade, genome-wide association studies (GWAS) have facilitated a multitude of discoveries in human genetics, identifying variants associated with a number of complex phenotypes and diseases^1,2^. This includes many neuropsychiatric traits, such as schizophrenia, for which over 100 genetic risk loci have been discovered through GWAS studies to date^3^. However, the journey from GWAS hit to biological function has proven challenging, for several reasons. Firstly, the sentinel SNP identified by GWAS is rarely the causal variant for the associated trait, but is instead a marker for a co-inherited genomic region known as the haplotype block^4^. The causal variant could lie anywhere within the haplotype block, which extends outwards to all SNPs that are in linkage disequilibrium (LD) with the GWAS SNP. Second, even if GWAS accurately pinpoints a variant, the mechanism by which that variant is causally associated with the trait in question can be uncertain^1^. A third and related challenge is that the majority of SNPs identified by GWAS fall outside of protein-coding parts of the genome, which has hindered their functional characterization^5-8^.

Several hypotheses exist to explain the preponderance of GWAS-identified SNPs in non-coding genomic regions. The most obvious is that these regions contain regulatory elements that have an influence on the trait in question. These may be *cis*-regulatory DNA sequences such as enhancers or silencers^9^. Alternatively, non-coding variants could affect chromatin looping, interfere with the binding of proteins or RNAs to DNA, or disrupt epigenetic marks^10^. These are all plausible, but another, not mutually exclusive, possibility is that these regions are transcribed into *cis-* and *trans-*acting non-coding (regulatory) RNAs, many of which may not yet be documented in public databases. Indeed, recent studies have indicated that human transcriptome annotations are far from complete, particularly in relation to long non-coding RNAs (lncRNAs)^11,12^.

lncRNAs are a diverse class of gene products that qualitatively constitute the major portion of the mammalian transcriptome, but which are still poorly catalogued and characterized^13^. The number of annotated lncRNAs has grown enormously in recent years, with almost 10,000 new lncRNA loci being added to GENCODE since 2009^11^, and many more likely to exist^14-17^. While only a small subset of annotated lncRNAs have been functionally characterized (e.g. *XIST, NEAT1, HOTAIR*), many have been shown to play unexpectedly diverse roles in epigenetic signalling and gene regulation including genetic imprinting, shaping chromosome conformation, forming subcellular organelles^18,19^, allosterically regulating enzymatic activity, and acting as scaffolds, guides, decoys or signals^20,21^. On average, lncRNAs are less abundant than protein-coding mRNAs, which can cause them to be missed (or dismissed as noise) by traditional RNA-seq assays^22^. This is due to the expression-dependent bias of RNA-seq, in which lowly expressed transcripts are less frequently sampled^23,24^. However, accumulating evidence shows that lncRNAs are not simply weakly expressed but rather are *precisely* expressed in highly specific patterns^25,26^. For example, a striking ∼40% of all lncRNAs are exclusively expressed in the brain^27^.

In order to address this problem, a technique known as RNA CaptureSeq has recently been developed. RNA CaptureSeq works by targeting specific genomic regions of interest for capture using oligonucleotide probes as baits, providing enhanced sequencing coverage of those regions^15,22,28^. RNA CaptureSeq has facilitated the identification of novel transcript isoforms within even well-studied loci such as *TP53*^*^15^*^, has provided the first genome-wide map of human splicing branchpoints^29^, and has proven particularly useful for the detection and quantification of lncRNAs and their many isoforms^14,22,30,31^. The increased sensitivity and resolution of RNA CaptureSeq make it a logical choice for the profiling of unannotated transcripts arising from non-coding GWAS haplotype blocks. To date, CaptureSeq has typically been performed using short-read RNA-seq, which relies on computational assembly of reads (∼100 bp) into transcript models. This process is notoriously difficult and error-prone, and does not provide certainty around isoform structures^32^. In contrast, long-read sequencing can sequence full-length transcripts ‘in one go’, and is able to resolve splicing events between distant exons^33^. However, long-read sequencing typically suffers from much lower throughput than short-read RNA-seq, and thus has almost exclusively been limited to profiling highly-expressed, protein-coding genes.

Here, we have used RNA CaptureSeq in conjunction with Oxford Nanopore Technologies (ONT) long-read cDNA sequencing in order to achieve an unprecedented level of sensitivity and resolution. While two recent studies have coupled CaptureSeq with Pacific Biosciences (PacBio) sequencing^14,30^, the present study represents the first use of CaptureSeq in conjunction with ONT sequencing to profile brain tissue. To investigate the transcriptional landscape of GWAS-identified genomic loci in human brain, we performed RNA CaptureSeq targeting 1,023 discrete haplotype blocks containing 1,352 non-coding GWAS SNPs associated with neuropsychiatric phenotypes. Transcripts were captured and sequenced from four regions of post-mortem brain tissue from three neurologically healthy donors. CaptureSeq was independently performed with both long-read and short-read RNA-seq, which provided orthogonal validation of our results. We used our recently developed set of spliced RNA spike-ins (‘sequins’)^23^ as internal controls to assess the efficiency of RNA capture, and also to benchmark the performance of ONT’s new PromethION instrument. We find that the majority (62.4%) of targeted haplotype blocks contain novel, multi-exonic transcripts, including 13% of targeted intergenic blocks. We uncover a wealth of unannotated transcripts, many of which are supported by independent evidence of transcription and show signs of potential functional roles. We present these transcriptomes as the foundation of an atlas of non-coding transcription in human brain that can be used to connect neuropsychiatric phenotypes with gene expression.

## RESULTS

### Selection of haplotype blocks and RNA CaptureSeq design

Using the NHGRI GWAS database^7,34^, we identified 1,023 discrete haplotype blocks across the genome containing 1,352 non-coding (intronic and intergenic) SNPs associated with neuropsychiatric phenotypes (see **Materials & Methods**). These phenotypes include behavioural traits, predisposition to addiction, mental illness and neurodegenerative disorders such as Alzheimer’s and Parkinson’s disease (see **Supplementary Table 1**). We designed tiling oligonucleotide probes targeting the selected GWAS-defined haplotype blocks, with annotated protein-coding exons (GENCODE^35^ v24) and repeat elements (RepeatMasker) omitted (**Fig. 1**). This resulted in a target territory of 96.2 Mb (∼3% of hg38). Of the 1,023 blocks targeted, 162 were intergenic (i.e. had no overlap on either strand with any GENCODE transcript). We employed targeted RNA capture (‘RNA CaptureSeq’)^28^ on RNA samples extracted from post-mortem brain tissue obtained from three neurologically healthy middle-aged males of European ancestry (see **Materials & Methods**). Brains were dissected into four different regions: caudate, prefrontal cortex (PFC), primary visual cortex (VCx) and superior colliculus (SupCol). Samples were spiked with RNA sequins^23^ (Mix A), a subset of which were also targeted for capture (25/78 genes; 49/164 isoforms) as part of our experimental design.

**Figure 1.**
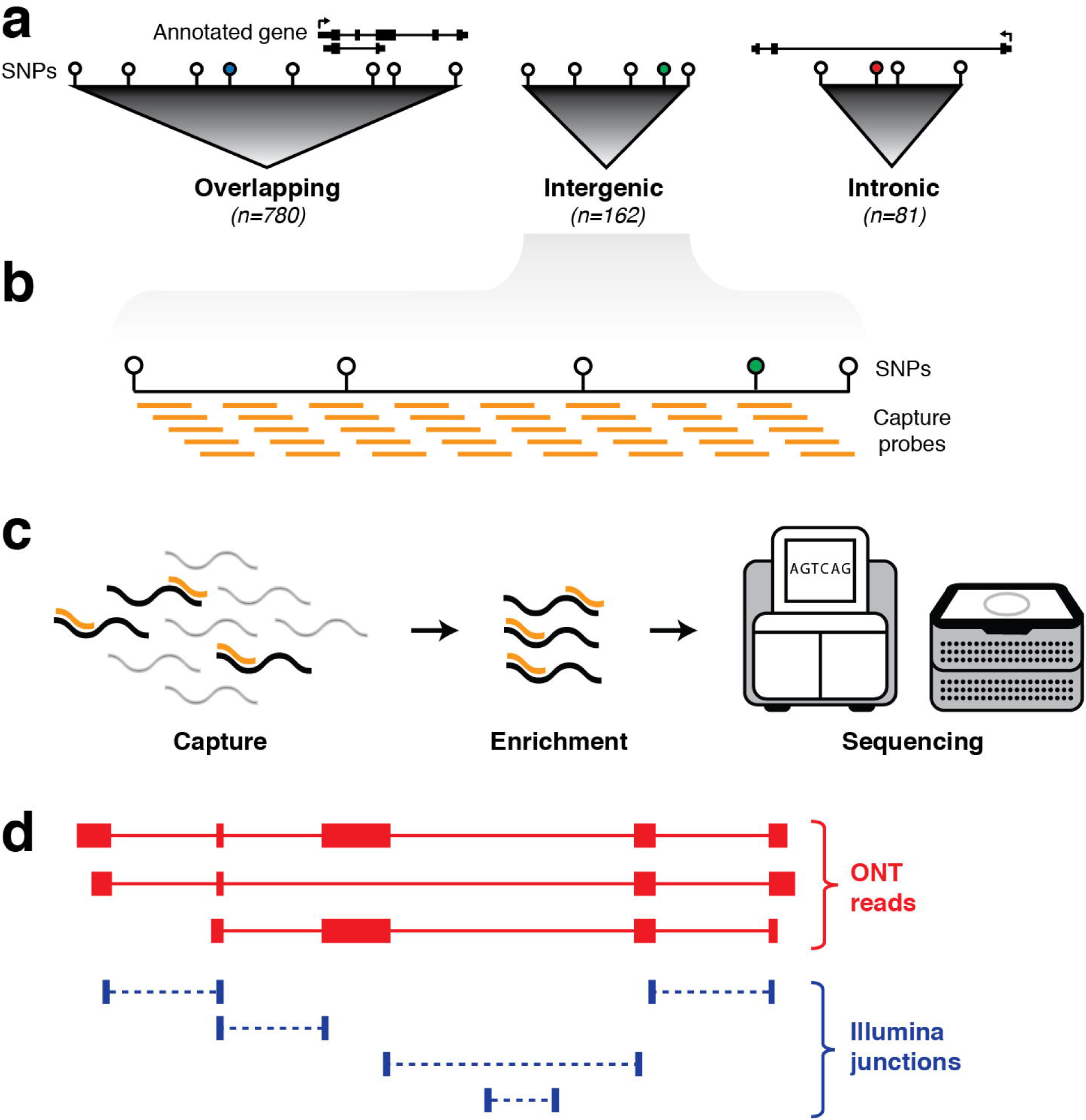
Schematic outline of experimental design. **(a)** First, haplotype blocks were predicted around 1,352 non-coding GWAS SNPs (coloured circles) associated with neurological phenotypes. Blocks were defined by identifying all SNPs in linkage disequilibrium (LD) with the GWAS SNP (white circles). Of the 1,023 blocks, 780 overlap an annotated exon (GENCODE v24), 81 are located entirely within an annotated intron, and 162 are intergenic. **(b)** Biotinylated oligonucleotide probes (orange bars) were designed to tile across haplotype blocks (with annotated protein-coding exons and repeat elements omitted). **(c)** Probes are used as baits to capture any transcripts generated from targeted regions, followed by pull-down enrichment and subsequent transcriptional profiling with both long- and short-read RNA sequencing. **(d)** Sequence reads are aligned to the genome (hg38) and a hybrid transcriptome is assembled by leveraging the advantages of both long- and short-read RNA-seq.

### Long-read transcriptional profiling using ONT cDNA sequencing

RNA CaptureSeq was performed as previously described^28^, with some minor modifications for the ONT libraries (see **Materials & Methods**). Following capture, we carried out single-molecule, long-read cDNA sequencing using ONT’s PromethION platform. This yielded a total of 62,553,766 base-called reads (58.7 Gbp of sequence) with a mean length of 1,035 nt and a median quality score of 8.9 (**Supplementary Fig. 1**). Reads were aligned using minimap2^36^ to a combined index comprising the human genome (hg38) and *in silico* chromosome (*chrIS*)^23^. Overall, we observed an alignment rate of 82.1%, with 56.9% of reads aligning to hg38 and 25.1% to *chrIS* (**Supplementary Fig. 2a**). We retained only reads with a perfect mapping score (mapQ=60). Of 22,967,553 reads that aligned to hg38 (mapQ=60), 13,187,936 mapped to regions targeted for capture, corresponding to an average on-target rate of 57.4% (**Supplementary Fig. 2c**). In parallel, we sequenced the same four brain cDNA samples using Illumina’s HiSeq 2500 instrument, yielding a total of 299,140,569 read pairs. Reads were aligned using STAR^37^ to the hg38 + *chrIS* reference. We observed an overall alignment rate of 78.4%, with 64.1% of reads aligning to hg38 and 14.3% to *chrIS* (**Supplementary Fig. 2b**). For the Illumina samples, we observed an average on-target rate of 83.4% (**Supplementary Fig. 2c**). By analyzing reads aligned to the *chrIS* reference sequence, we calculated an overall error rate for the PromethION of 10.8733% (mismatch rate 5.11%; indel rate 5.7633%) (**Supplementary Fig. 2d**). This was ∼50-fold higher than the rate observed for Illumina reads, which had an overall error rate of 0.2341% (mismatch 0.2241%; indel 0.01%).

**Figure 2.**
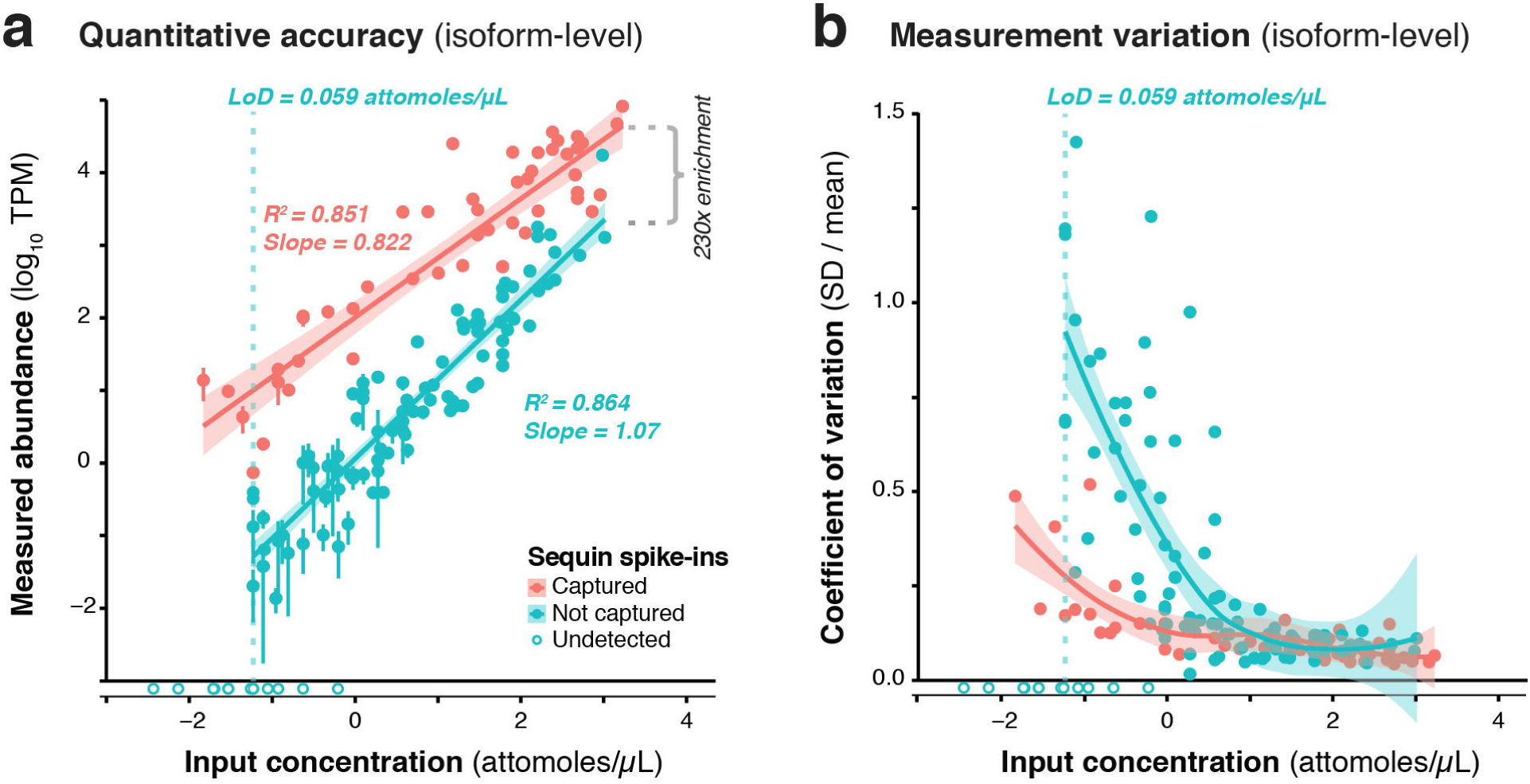
Validation of CaptureSeq design and ONT sequencing. **(a)** A subset of RNA sequins were targeted for capture as part of our CaptureSeq design (*n*=25/78 genes; 49/164 isoforms). By plotting the measured abundance (TPM; y-axis) against the known input concentration (x-axis) for captured (red) and non-captured (green) sequin isoforms, we can compare the quantitative accuracy of captured vs non-captured transcripts. Open circles indicate sequin isoforms that were not detected. Vertical dotted green line indicates the limit of detection (LoD) for non-captured transcripts. By comparing the difference between the measured abundance of captured and non-captured transcripts, we observe a ∼230-fold enrichment of CaptureSeq. Error bars represent standard deviation (SD) between the four replicate ONT samples. **(b)** Scatter plot shows the coefficient of variation (SD divided by mean) of each spike-in plotted against its respective input concentration, indicating the expression dependent bias of RNA-seq.

### Validation of RNA CaptureSeq design and quantitative accuracy

We used RNA sequins to assess the efficiency of RNA capture and the quantitative accuracy of ONT sequencing using the PromethION device. We carried out isoform-level quantification using Salmon^38^, finding a strong correlation between measured abundance (transcripts per million; TPM) and input concentration for captured (R^2^=0.851, slope=0.822) and non-captured (R^2^=0.864, slope=1.07) sequins alike (**Fig. 2a**). By calculating the average difference between captured and non-captured sequins at each matched concentration point, we observed a ∼230-fold enrichment of CaptureSeq (**Fig. 2a**). At higher concentrations, we observed diminishing capture efficiency, which corresponds to saturation of capture probes but is unlikely to affect transcripts within the physiological range of gene expression^22,28^. While we successfully detected all captured sequins, we failed to detect the 10 sequin isoforms of lowest input concentration that were not targeted for capture, equating to a lower limit of detection (LoD) of 0.059 attomoles/µL. Further, to assess the variation of expression measurements at different concentrations, we plotted the coefficient of variation (CV; SD divided by mean) for each sequin against its input concentration, observing that CV declined with increasing concentration for both captured and non-captured standards (**Fig. 2b**). This illustrates the expression dependent bias of RNA-seq, whereby lowly expressed transcripts are less accurately quantified. We also quantified sequins at the gene-level (reads per gene per 10k reads; RPG10K^39^), observing similar results (**Supplementary Fig. 3**).

**Figure 3.**
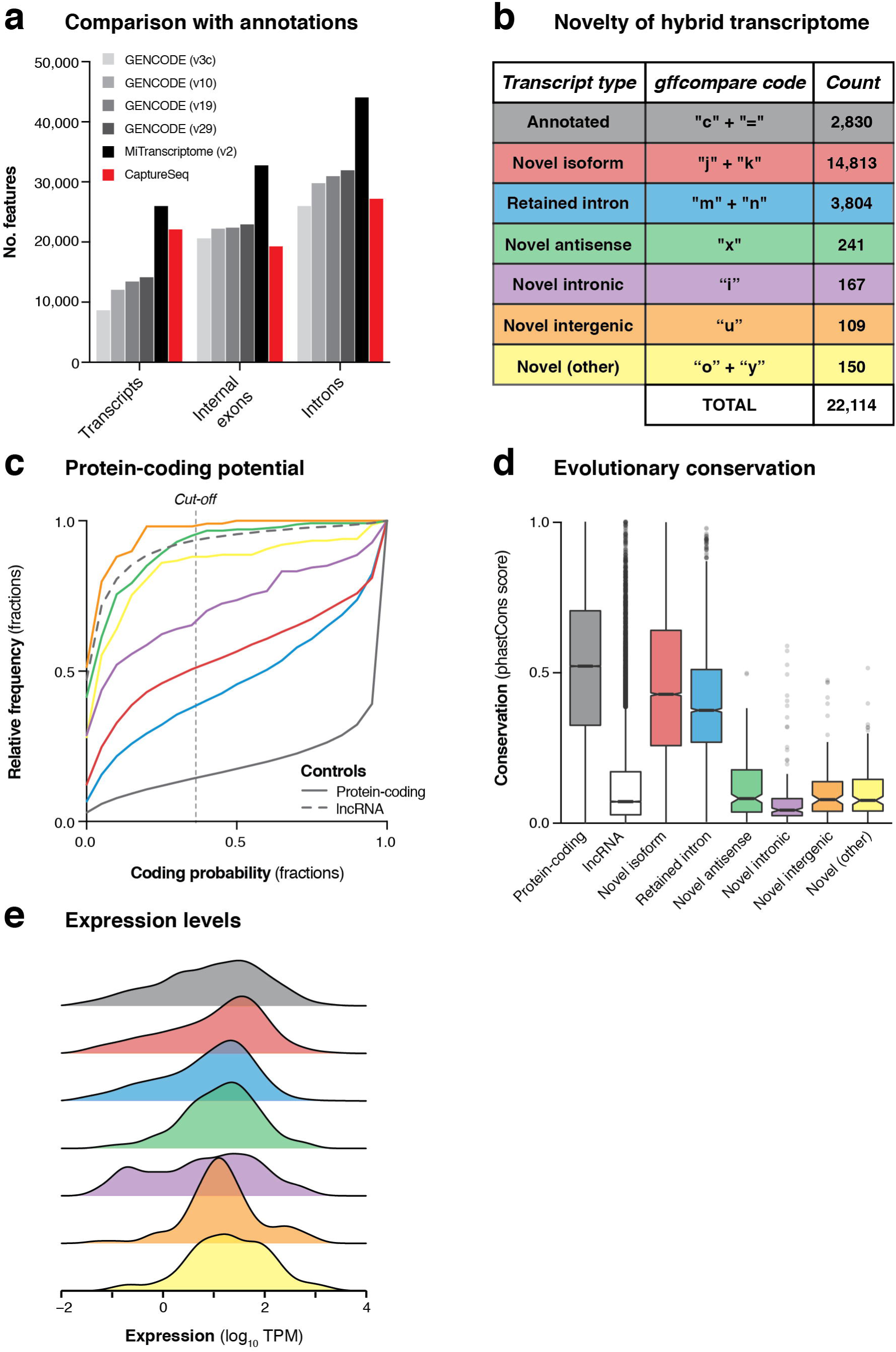
Transcriptional landscape of haplotype blocks associated with neuropsychiatric functions. **(a)** Bar charts show the number of transcripts, introns and internal exons contained in our filtered hybrid transcriptome (red) compared to existing annotations (four versions of GENCODE and MiTranscriptome v2). Only multi-exonic transcripts that overlap with targeted haplotype blocks are considered in this analysis. **(b)** Table shows the classification of transcripts in our hybrid transcriptome in relation to the latest GENCODE annotation (v29). **(c)** Cumulative frequency histograms show the coding potential of our transcripts, as assessed by CPAT. Colours refer to the categories defined in part (b). Vertical dotted line indicates the commonly used cut-off for human transcripts (0.364). GENCODE (v29) annotated lncRNAs (grey dotted line) and protein-coding genes (grey solid line) are also plotted for reference. **(d)** Box plots show the distribution of phastCons scores^75^ (vertebrate 100-way alignment) of our transcripts, coloured by type. Box edges indicate lower and upper quartiles, centre lines indicate median, notches indicate the 95% confidence interval around the median. **(e)** Density plots show the mean expression (log_10_ TPM) of transcripts across all four samples, as measured by Illumina short-read sequencing.

### Concordance of splice junction detection between ONT and Illumina

Our spliced ONT reads aligned to hg38 (mapQ=60) contained a total of 939,558 splice junctions. In comparison, we detected a total of 234,138 uniquely mapped junctions in the matched Illumina data. We found that 164,183 of these junctions were shared between the two technologies; that is, 17.5% of junctions detected using ONT were validated by Illumina, and 70.1% of junctions detected by Illumina were validated by ONT (**Supplementary Fig. 4a**). Of the ‘shared’ junctions, 144,227 (87.8%) were previously annotated in GENCODE (v29), while the remaining 19,956 junctions were novel (12.2%) (**Supplementary Fig. 4b**). ONT detected an additional 32,777 GENCODE junctions that Illumina missed, while Illumina detected an additional 18,860 GENCODE junctions that ONT missed (**Supplementary Fig. 4c**). Of the 19,956 ‘shared novel’ junctions, we found that 2,036 (10.2%) were validated by two recent studies which coupled RNA CaptureSeq with Pacific Biosciences (PacBio) long-read sequencing^14,30^, providing independent evidence for their credibility.

**Figure 4.**
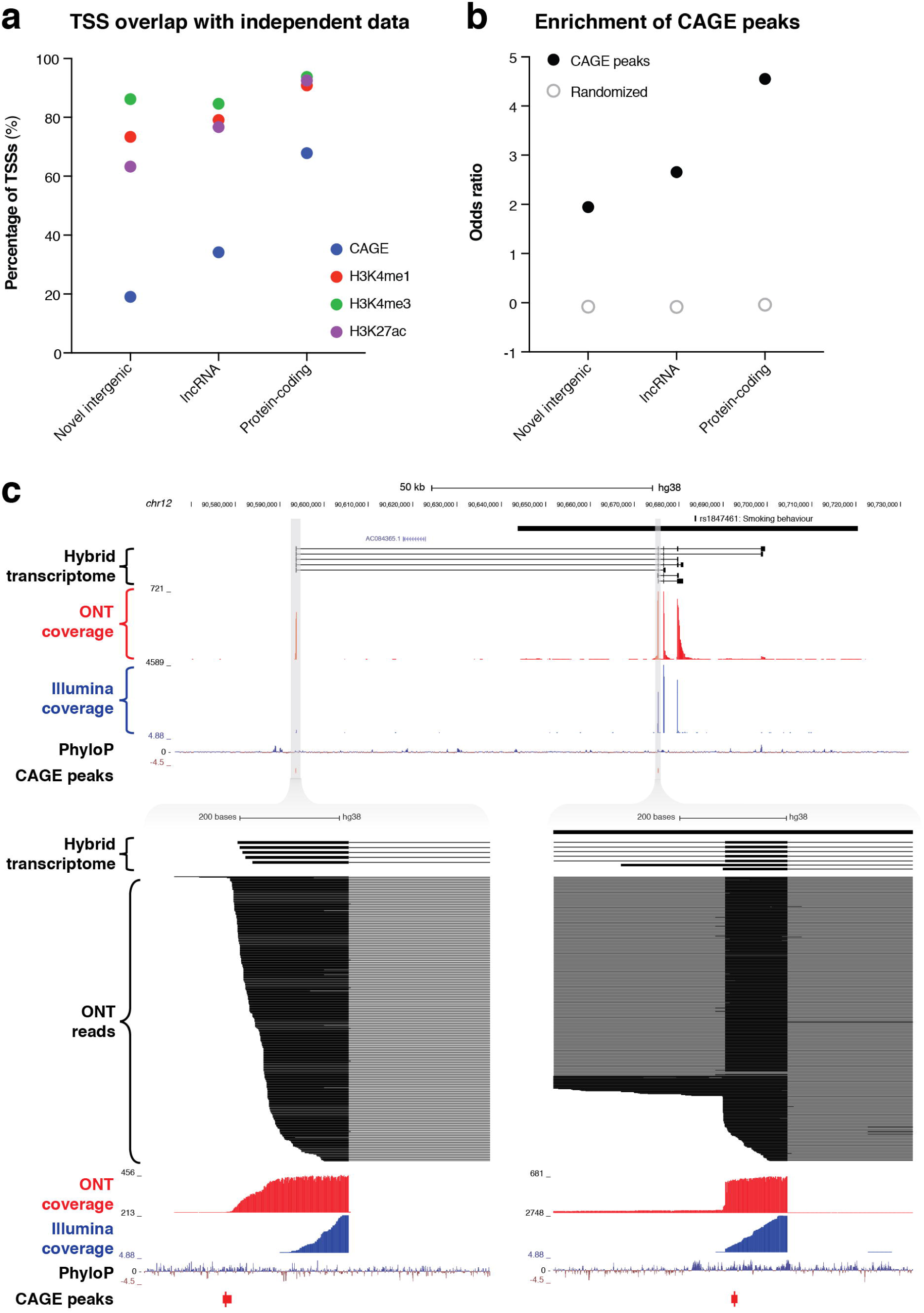
Identification of novel intergenic transcripts. **(a)** Bar charts indicate the fraction of transcription start sites (TSSs) occupied by cap analysis of gene expression (CAGE) peaks (blue), as well as the three epigenetic marks that are typically associated with actively transcribed promoters: H3K4me1 (red), H3K4me3 (green) and H3K27ac (purple). GENCODE (v29) lncRNAs and protein-coding genes are also plotted for reference. **(b)** Bar charts show the enrichment of the promoter regions of transcripts for CAGE peaks. Odds ratio of enrichment is plotted for novel intergenic transcripts compared to lncRNAs and protein-coding genes. **(c, upper)** Genome browser view shows a novel intergenic locus identified overlapping a GWAS haplotype block (solid black bar, top) associated with smoking behaviour (rs1847461) on chromosome 12. Transcripts from our filtered hybrid transcriptome are shown below, followed by spliced ONT sequencing coverage (red), spliced Illumina sequencing coverage (blue), PhyloP conservation track, and CAGE robust peaks. **(c, lower)** Two separate magnified views show novel TSSs supported by CAGE peaks and highly-conserved promoter regions. ONT read alignments are also shown.

### Benchmarking the performance of ONT cDNA sequencing

We next used RNA sequins to assess the performance of the PromethION instrument in accurately sequencing full-length transcripts. In contrast to the ERCC spike-ins, sequins have a more realistic range of transcript sizes, with 15 isoforms ≥2.5 kb in length. Of these 15, only six were fully sequenced ‘in one go’ with a single ONT read. The longest sequin isoform that was fully sequenced with single reads was *R2*_*26*_*1*, with a 4,375 nt read that covered all 18 of its exons (**Supplementary Fig. 5a**). The longest read that mapped to *chrIS* was 5,213 nt, which represented the majority of the *R2*_*19*_*2* isoform (6.9 kb in total). Another notable example was a 3,473 nt read which mapped to *R1*_*24*_*1* (4.6 kb in total), spanning 33 out of 36 exons. Most synthetic isoforms <2.5 kb were fully sequenced with single ONT reads, allowing complex, alternatively spliced synthetic loci to be unambiguously deconvoluted (**Supplementary Fig. 5b**). We observed a similar distribution of mapped reads lengths aligning to *chrIS* (mean=762 nt) and hg38 (mean=937 nt) (**Supplementary Fig. 2e**).

**Figure 5.**
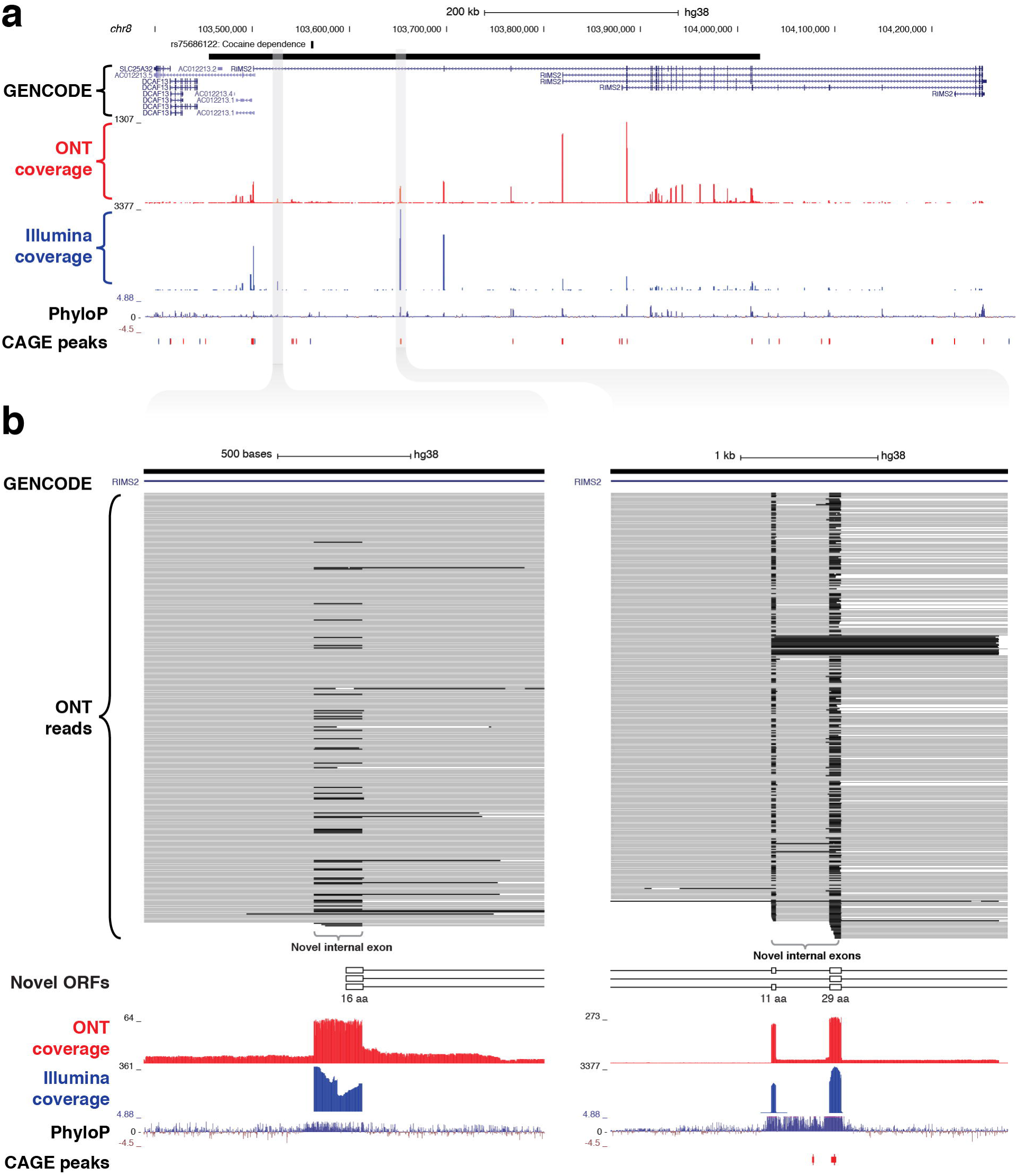
Identification of novel internal coding exons in *RIMS2* gene. **(a)** Genome browser view shows a ∼570 kb GWAS haplotype block (solid black bar, top) on chromosome 8 associated with cocaine dependence (rs75688122). CaptureSeq detected multiple novel splice isoforms of *RIMS2*, a gene involved in neurotransmitter release. **(b)** Magnified views show three novel, highly conserved exons detected in the first intron of *RIMS2*, which are predicted to collectively add 56 amino acids to the start of the RIMS2 protein. The first novel internal exon includes a start codon (left), while the second and third exons (right) are in-frame (33 and 87 bp, respectively).

We used the gffcompare tool^40^ to assess the sensitivity and precision of our synthetic transcriptome in relation to the *chrIS* annotation, at the base-, exon-, intron-, transcript- and gene-levels. When considering the raw read alignments, we observed high sensitivity across these features, but large numbers of false-positive events led to poor precision scores (**Supplementary Fig. 6a**). This is due to the higher error rate of ONT sequencing, which leads to spurious alignments around the boundaries of exon junctions (**Supplementary Fig. 5c**). We compared the number of reads spanning annotated *chrIS* junctions (true-positives) and unannotated *chrIS* junctions (false-positives), finding that ONT performed poorly in comparison to Illumina (**Supplementary Fig. 6b**). We used the ‘Pinfish’ suite of tools (https://github.com/nanoporetech/pinfish) to cluster together ONT reads (mapQ=60) having similar exon/intron structures, and then create a consensus of the clusters by calculating the median of exon boundaries from all transcripts in the cluster (see **Materials & Methods**). After clustering, we observed marked improvements in precision with only minor drops in sensitivity, validating that the approach was effective in processing raw ONT reads (**Supplementary Fig. 6c**). After benchmarking the ‘Pinfish’ clustering approach using RNA sequins, we went on to apply this to ONT reads mapping to hg38 (mapQ=60). This generated a set of 10,528 consensus multi-exonic transcripts that overlapped targeted regions, comprising 12,369 unique splice junctions. We found that the rate of ONT junction validation by Illumina data increased from only 17.5% before clustering to 83.0% after clustering (**Supplementary Fig. 6d**).

Using sequins, we observed that ONT sequencing had a slight length bias, with synthetic transcripts at around the ∼1 kb size over-represented and transcripts at either end of the size spectrum under-represented. This effect was seen for both captured (**Supplementary Fig. 7a**) and non-captured (**Supplementary Fig. 7c**) sequins, suggesting that the length bias was caused by ONT sequencing rather than by the process of capture. Similarly, we observed a slight GC content bias for captured sequins (**Supplementary Fig. 7b**), but this effect largely disappeared for non-captured sequins (**Supplementary Fig. 7d**), implying that GC bias arose due to the process of capture rather than ONT sequencing. In most cases, ONT sequencing had more uniform coverage of exons compared to Illumina (**Supplementary Fig. 8a,b**). With Illumina sequencing, many exons exhibited highly uneven coverage profiles that were reproducible between replicates (**Supplementary Fig. 8a,b**). We calculated the coverage of each base in every sequin exon targeted for capture (n=279), finding that there was significantly less variation between bases for ONT (median CV = 0.0339) than Illumina sequencing (median CV = 0.172) (**Supplementary Fig. 8c**). This difference was statistically significant (paired t-test; p-val < 0.0001).

### Defining a hybrid transcriptome

Long-read sequencing provides accurate full-length isoform structures, but suffers from a relatively high sequencing error rate that leads to spurious splice junction detection^41^. Conversely, short-read sequencing relies on the error-prone computational reconstruction of short reads into transcript models, but has highly accurate splice junction detection^32^. We sought to leverage the advantages of both technologies by defining a hybrid transcriptome that incorporates ONT reads with splice junctions corrected using matched Illumina short-read data. To do so we used the FLAIR^41^ tool, which corrects misaligned ONT splice junctions using genome annotations and accompanying Illumina junctions (see **Materials & Methods**). Of 3,586,593 spliced ONT reads that mapped to hg38, we were able to successfully correct 2,422,358 of these (67.5%) using GENCODE annotations (v29) and our matched Illumina splice junction data. These corrected reads were then collapsed into a non-redundant transcriptome.

Next, we developed a comprehensive pipeline to retain only high-confidence transcripts that overlapped targeted haplotype blocks (see **Materials & Methods**). We filtered out all transcripts that met any of the following criteria: (i) no overlap with any capture probe, (ii) contained <3 exons; (iii) contained an intron >1 Mb, (iv) contained a non-canonical splice junction (i.e. not GT-AG, GC-AG or AT-AC); (v) had any of the following gffcompare^40^ classification codes: ‘e’, ‘p’, ‘r’, ‘s’; (vi) was <200 nt in length; or (vii) was redundant or contained wholly within another transcript. These filtering steps produced a set of 22,114 multi-exonic transcripts overlapping GWAS haplotype blocks associated with neuropsychiatric functions. These transcripts collectively comprised 27,181 unique introns (splice junctions) and 19,288 discrete internal exons (**Fig. 3a**). This represents a comparable number of features as are contained in the GENCODE (v29) annotation across the genomic regions we targeted for capture (**Fig. 3a**). While our transcriptome was less comprehensive than the MiTranscriptome annotation (v2)^42^, the latter was generated by merging ∼6,500 independent RNA-seq datasets into a consensus set, while our dataset was generated in just two experiments using four brain samples.

### Transcriptional landscape of haplotype blocks associated with neuropsychiatric functions

We compared our hybrid transcriptome to the GENCODE (v29) annotation using gffcompare^40^. Only 2,830 transcripts (12.8%) were exact matches of (or contained within) GENCODE transcripts, while the remainder were categorized as putative novel transcripts (**Fig. 3b**). Most of the novel transcripts represented unannotated splice isoforms of known genes, but we also detected 241 novel antisense transcripts (no GENCODE overlap on the same strand) and 109 novel intergenic transcripts (no GENCODE overlap on either strand) (**Fig. 3b**). We assessed the protein-coding capacity (**Fig. 3c**) and evolutionary conservation (**Fig. 3d**) of our transcriptome, finding that the novel antisense and novel intergenic transcripts closely resembled annotated lncRNAs. Overall, we found that 638/1,023 (62.4%) targeted haplotype blocks contained at least one novel transcript, including 21/162 (13.0%) targeted intergenic blocks. We used the matched short-read RNA-seq data to quantify the expression of our hybrid transcriptome (**Fig. 3e**). We also assessed the concordance between ONT and Illumina quantitative expression measurements for our transcriptome. Using Salmon^38^ with our hybrid transcriptome as a reference, we quantified the expression of transcripts in each sample individually, observing reasonably strong concordance between the two orthogonal technologies (Spearman correlation coefficient (ρ) for SupCol: 0.612; PFC: 0.616; VCx: 0.650; caudate: 0.615) (**Supplementary Fig. 9**).

### Analysis of novel intergenic transcripts

The combined resolution of CaptureSeq with ONT long-read sequencing enabled us to identify 109 novel intergenic transcripts overlapping GWAS SNPs associated with neuropsychiatric traits. Only two of these transcripts were predicted to have protein-coding potential; the remaining 107 are therefore classed as putative lncRNAs (**Fig. 3c**). We investigated whether any of the novel intergenic transcripts were supported by independent signatures of transcription, such as cap analysis of gene expression (CAGE) peaks or epigenetic marks. We found that 21 transcripts (19.3%) had an annotated FANTOM CAGE robust peak^43^ in the vicinity of their TSS (within 500 bp on the same strand) (**Fig. 4a**). This represented a significant enrichment (odds ratio = 1.95) compared to randomized regions, but did not reach the enrichment of CAGE peaks in the TSSs of annotated lncRNAs (odds ratio = 2.65) or protein-coding transcripts (odds ratio = 4.55) (**Fig. 4b**). Notably, of our intergenic transcripts that were not supported by CAGE peaks, 8/88 (9.1%) were independently validated by a recent study that coupled CaptureSeq with PacBio long-read sequencing^30^, thus providing orthogonal evidence for their veracity (**Supplementary Fig. 10**). In several of these examples, our hybrid assembly expanded upon the PacBio assembly, identifying additional exons and splicing events. Further, the three epigenetic marks that are typically associated with actively transcribed promoters (H3K4me1, H3K4me3 and H3K27ac) were present near the TSS of 80 (73.4%), 94 (86.2%) and 69 (63.3%) of the novel intergenic transcripts, respectively (Roadmap Epigenomics Consortium^44^) (**Fig. 4a**). Finally, we overlapped our novel intergenic transcripts with a recent dataset of conserved RNA structures (CRSs)^45^. We found that 9/109 transcripts (9.2%) contained one or more CRSs overlapping their exons, which may be indicative of RNA-mediated functionality.

For example, we discovered a novel intergenic locus overlapping a GWAS haplotype block associated with ‘smoking behaviour’ (rs1847461)^46^ on chromosome 12 (**Fig. 4c, upper**). We identified 7 multi-exonic transcripts at this locus, with all splice junctions confirmed by Illumina short-read sequencing. Many exons were highly conserved, and the two TSSs were supported by CAGE peaks (**Fig. 4c, lower**). Another example included a novel intergenic locus overlapping a GWAS SNP associated with multiple sclerosis (rs354033)^47^ on chromosome 7 (**Supplementary Fig. 11**). Once again, splice junctions were validated with Illumina sequencing, several exons were highly conserved, and most transcripts overlapped a CAGE peak at the 5’ end.

### Identification of novel isoforms in well-studied genes with neuropsychiatric functions

As well as discovering novel gene loci in the vicinity of GWAS neuropsychiatric SNPs, CaptureSeq enabled us to identify hundreds of novel splicing events in annotated genes with established roles in neuropsychiatric function. For example, we captured a ∼570 kb haplotype block on chromosome 8 that includes a SNP associated with cocaine dependence (rs75686122)^48^ (**Fig. 5a**). This SNP is located within the first intron of the *RIMS2* gene, which encodes a protein that interacts with various synaptic proteins that are important for normal neurotransmitter release^49^. We identified three novel internal exons in the first intron of *RIMS2* that are highly conserved and are predicted to encode novel ORFs (**Fig. 5b**). The first novel exon encodes a start codon and the first 16 amino acids of the novel ORF, while the second and third exons are in-frame and encode another 11 and 29 amino acids, respectively. Collectively, these novel exons are predicted to add 56 amino acids to the start of the RIMS2 protein. Whilst interesting, this finding would require further proteomic validation.

In another example, we captured a ∼4 kb haplotype block containing a GWAS SNP associated with schizophrenia (rs12807809)^50^, located immediately upstream of the *NRGN* gene on chromosome 11 (**Supplementary Fig. 12**). *NRGN* encodes a postsynaptic protein that is thought to be a direct target for thyroid hormone in human brain^51^. We identified a novel TSS located ∼20 kb upstream of the annotated TSS for *NRGN* that was supported by a CAGE peak and had a highly conserved promoter region. The novel transcripts incorporate annotated exons of *NRGN*, such that the novel introns span SNP rs12807809.

### Using ONT sequencing to detect coordination between distant exon pairs

One of the promised advantages of long-read sequencing is that it can deconvolute coordinated alternative splicing events between distant pairs of exons^33,52^. This type of information cannot be gleaned using short-read RNA-seq because distant exon pairs are never sequenced on the same fragment. Since several RNA sequin loci provide known examples of coordinated splicing between distant exon pairs, we first used *chrIS* as a proof-of-principle to verify that ONT sequencing could reliably detect such events. For example, the sequin locus *R1*_*22* comprises two alternatively spliced isoforms that contain a distant mutually associated pair (dMAP) of exons (**Supplementary Fig. 13a**). With ONT sequencing, we were able to accurately resolve the long-range connectivity between these exons. However, with short-read Illumina sequencing, computational assembly (with StringTie^53^) produced a false-positive transcript structure in which one exon in the pair was included and the other excluded. Likewise, the *R1*_*103* sequin locus comprises two alternative isoforms that contain a distant mutually exclusive pair (dMEP) of exons (**Supplementary Fig. 13b**). Again, ONT sequencing enabled their long-range relationship to be resolved, while short-read sequencing produced a false-positive transcript structure in which both exons were present. We used our previously published methods^33,54^ to search for similar examples of coordinated alternative exon pairing in the human genome. For example, we detected a dMEP of exons in *MBNL2*, a gene implicated in the development of myotonic dystrophy that overlaps a targeted haplotype block associated with alcoholism (rs9556711)^55^ on chromosome 13 (**Supplementary Fig. 13c**). This exon pair (exons number 7 and 9 of transcript ENST00000469707.5) was never simultaneously present on the same molecule, with no reads containing both exons, 19 reads containing only exon 7 but not exon 9, 37 reads containing exon 9 but not exon 7, and 50 reads which skipped both exons (**Supplementary Fig. 13c**). This coordination was highly significant, with a Fisher’s exact test p-value of 1.23 × 10^−4^.

## DISCUSSION

This study has successfully employed RNA CaptureSeq to reveal the rich transcriptional diversity hidden within non-coding regions of the genome that have previously been associated by GWAS with neuropsychiatric functions. In doing so, we have vastly expanded existing transcriptome annotations of these regions, creating an expression atlas of thousands of novel isoforms in human brain. Many of these assembled isoforms show preliminary evidence of functional roles, including highly conserved exons and promoter regions, TSSs supported by CAGE peaks and epigenetic marks, and novel predicted ORFs.

The improved sensitivity of CaptureSeq allowed us to assemble 109 novel intergenic transcripts in regions of the genome that were previously thought to be transcriptionally silent. As a class, these transcripts resemble annotated lncRNAs, as judged by their protein-coding probability scores, relatively low overall expression and evolutionary conservation compared to protein-coding genes, which is typical of regulatory sequences^56^. While only a minority (∼19%) of these were supported by CAGE peaks, their TSSs were nonetheless enriched for CAGE peaks compared to the genomic background. It is also worth pointing out that CAGE data is itself expression-dependent, and the fact that ∼9% of our novel intergenic transcripts lacking CAGE support were independently validated by a recent study^30^ implies that the sensitivity of CaptureSeq surpasses that of CAGE over targeted regions.

To date, a poor understanding of the sequence-function relationship of lncRNAs (as opposed to protein-coding genes) has hindered their functional characterization. Promisingly, several technologies for probing lncRNA functions and mechanisms have begun to emerge, including ChIRP-seq (for assaying DNA/protein binding partners) and SHAPE-seq (for RNA structure)^57^. Further, two landmark studies have recently employed CRISPR-Cas9 genome editing to perturb lncRNA loci *in vivo*, leading to the identification of hundreds of lncRNAs that have an effect on cell growth^58,59^. Mechanistically, non-coding GWAS SNPs could be located within the promoters of lncRNAs, hence influencing their expression. Alternatively, they could fall within lncRNA exons, thereby potentially affecting RNA secondary structure (so-called ‘riboSNitches’)^60,61^. Indeed, we found that ∼9% of our novel intergenic transcripts had exonic overlap with one or more CRSs^45^, providing potential evidence for RNA-mediated functionality. GWAS SNPs located in proximity to exon boundaries can potentially alter splicing patterns in these loci, which display complex transcriptional activity.

It is conceivable that some of our novel intergenic transcripts represent the output of enhancers. Indeed, recent evidence indicates that most – if not all – enhancers are transcribed into non-coding RNAs that have been termed ‘eRNAs’^62^. However, there is still significant controversy around whether eRNA transcripts are functional *per se*, or whether it is merely the act of their transcription that is indicative of some underlying function^62^. Of the intergenic haplotype blocks we targeted for capture, we failed to detect transcription in the majority of cases. However, it must be borne in mind that we only polled four brain regions and that many non-coding RNAs are only expressed in very specific cell populations^26,63^.

This study also demonstrated the utility of spike-in controls in analyzing NGS data^64^. The use of RNA sequins^23^ provided a faithful set of internal controls that helped to validate our experimental design, including an assessment of capture efficiency, sensitivity and quantitative accuracy. Of the 15 sequin isoforms ≥2.5 kb in length, six were fully sequenced ‘in one go’, including a ∼4.4 kb synthetic transcript comprising 18 exons. Furthermore, we showed how the *in silico* chromosome *(chrIS)* can act as a comprehensive ‘ground truth’ reference against which FP and FN findings with RNA-seq can be evaluated. Because *chrIS* emulates the features of a real human chromosome, we would expect similar rates of TP and FP events for human transcripts. Notably, this type of analysis is not possible with previous spike-in controls (e.g. the ERCC spike-ins^24^), because they are mono-exonic and do not recapitulate the complexity of eukaryotic gene splicing. Despite these advantages, this study also illustrates some of the limitations of spike-in controls. While sequins were added to samples at a low fractional concentration (2%), we found that a disproportionally high fraction of reads aligned to *chrIS*, thereby sacrificing reads that could otherwise have come from endogenous transcripts. While we added sequins in proportion to each sample, the process of capture positively selects on the targeted genomic regions in a way that is not entirely predictable, thereby inadvertently changing the ratio of sample to spike-ins.

In conclusion, this study substantially expands the transcriptome annotations for regions of the genome associated with important neuropsychiatric traits, including diseases like Alzheimer’s, Parkinson’s and schizophrenia. These transcriptomes collectively comprise a valuable atlas that can be used to connect gene expression with neuropsychiatric traits. Ultimately, novel transcripts identified herein could act as biomarkers for disease or potential therapeutic targets.

## MATERIALS & METHODS

### Selection of GWAS haplotype blocks

The complete GWAS database was downloaded from the NHGRI catalog^7,34^, then filtered to include only studies with a sample size of n ≥1000 that focused on traits associated with the brain (including behavioural traits, mental illness, as well as neuropsychiatric disorders like Alzheimer’s, Parkinson’s and schizophrenia). Since GWAS simply associates traits with regions of the genome that are in linkage disequilibrium (LD), the causative SNP could be anywhere within the region of LD. In order to ensure the causative SNP is included, it is necessary to capture the full haplotype block associated with the SNP reported by GWAS. As such, we used SNAP^65^ to identify all SNPs in LD, with an LD threshold of R >0.5 and a maximum distance between SNPs of 500 kb. All SNPs in LD were then assumed to comprise a single haplotype block. This resulted in 1,323 haplotype blocks comprising a total of 1,352 GWAS SNPs associated with neuropsychiatric phenotypes (see **Supplementary Table 1**). Since some of these haplotype blocks overlapped, we merged them into a set of 1,023 discrete, non-overlapping blocks.

### Probe design

Biotinylated oligonucleotide probes were designed to tile across the abovementioned 1,023 blocks, excluding any protein-coding exons (GENCODE v24) and repeat elements (RepeatMasker). Probes were designed in accordance with previous guidelines^28^. This resulted in a final capture space of 96,234,476 bp. In addition, a subset of our RNA sequins spike-ins^23^ were targeted for capture (25/78 genes; 49/164 isoforms). In selecting spike-ins to target, we chose transcripts that spanned across the range of concentrations in the staggered mixture (Mix A). Probe designs were submitted to Roche NimbleGen for synthesis.

### Human brain samples and RNA extraction

Brain tissue from three de-identified, neurologically healthy males of European ancestry was supplied by the NSW Brain Tissue Resource Centre (BTRC) under Project ID 0379 and ethics approval number HC16189. Donors were aged between 61 and 64, had cardiac causes of death, with post-mortem intervals of brain collection between 17-41 h. From each brain, 3 mg of tissue was dissected by the BTRC from four regions: prefrontal cortex (PFC), primary visual cortex (VCx), caudate, and superior colliculus (SupCol). RNA was extracted from samples using the protocol for QIAGEN’s miRNeasy kit: 700 µL QIAzol was added per sample on ice, which were then transferred to 2 mL tubes containing a single 5 mm ball bearing and lysed on the Tissue Lyser II for 2 minutes at 20 Hz. Upon completion of the cycle, adapters were rotated and shaken for an additional 2 minutes at 20 Hz. Once lysed, the samples were purified according to the miRNeasy mini kit protocol instructions, including the on-column DNAse treatment. Elution was in 15 µL nuclease-free H_2_O, with the eluate being placed back onto the columns for an additional run in order to ensure maximum recovery and concentration.

### cDNA generation for Oxford Nanopore Technologies (ONT) sequencing

RNA CaptureSeq was performed as previously described^28^, with the following modifications for the ONT platform. The samples were prepared by combining 1 µg of each of the four human brain samples (SupCol, PFC, VCx, and caudate) independently with 10 ng of RNA sequins^23^ (Mix A; 2% fractional abundance). Volumes were adjusted to 11.5 µl per sample with nuclease-free water, followed by the addition of 0.5 µl of 250 mM Random Primer 6 (S1230S, NEB) and 1 µl Deoxynucleotide Solution Mix (N0447, NEB). Samples were mixed by tapping, briefly spun by microfuge, incubated at 65°C for 5 min and immediately cooled on ice. 4 µl of First Strand buffer and 2 µl 100mM DTT (SuperScript II Reverse Transcriptase: 18064014, ThermoFisher) were added to the samples, which were mixed by tapping, briefly spun by microfuge and incubated at 42°C for 2 min. For the reverse transcription reaction, 1 µl of SuperScript II Reverse Transcriptase was added to the 19 µl reaction mixes, which were mixed by tapping, briefly spun by microfuge and incubated with the following thermocycler conditions: RT reaction at 50°C for 50 min; denaturation at 70°C for 15 min; and holding indefinitely at 4°C. The second strand synthesis reaction mix was assembled as follows (NEBNext Ultra II Non-Directional RNA Second Strand Synthesis Module: E6111S, NEB): 20 µl of first strand cDNA, 10 µl buffer, 5 µl enzyme mix, and 45 µl nuclease-free water. Reactions were incubated with the following thermocycler conditions: second strand synthesis at 16°C for 1 h; and holding indefinitely at 4°C. Resulting cDNA was purified by a 10 min incubation with 144 µl (1.8X) Agencourt AMPure XP beads (A63880, Beckman Coulter), 2x 200 µl washes with 80% ethanol for 30 sec each, 5 min air drying, and elution in 52 µl nuclease-free water.

Following reverse transcription, the four samples had an average fragment length of 1500 bp with the following concentrations: 6.99 ng/µl (SupCol), 5.56 ng/µl (PFC), 5.5 ng/µl (VCx), and 6.5 ng/µl (caudate) according to a genomic DNA screentape analysis (Agilent 5067-5365, 5067-5366). The samples were each end-prepped and dA-tailed using the NEBNext Ultra II End Repair/dA-Tailing Module (E7595) as follows: 45 µl DNA, 7 µl Ultra II End-prep reaction buffer, 3 µl Ultra end-prep enzyme mix, and 5 µl nuclease-free water. Each reaction was incubated at 20°C for 30 min followed by 65°C for 30 min. Following end-prep, the samples were purified using AMPure beads (A63882) at a 1:1 ratio with incubation at room temperature for five minutes, two 140 µl washes with 70% ethanol, and elution for two minutes in 31 µl nuclease-free water. The PCA adapters from the ONT ligation sequencing kit 1D (SQK-LSK108) were ligated onto the end-prepped samples as follows: 30 µl of the end-prepped DNA, 20 µl PCR adapters (PCA), and 50 µl Blunt/TA ligase master mix (NEB) were combined and incubated at room temperature for 10 minutes. The adapted DNA was purified using Agencourt AMPure Beads as described above. The DNA was then eluted in nuclease-free water. Next, each sample was amplified in a 100 µl reaction using NEB Long-Amp Taq (M0323) as follows: 1X LongAmp Taq reaction buffer, 300 µM dNTPs, 2 µl PRM primers, and 48 µl DNA from the previous step. The reaction for each sample was divided into two 50 µl reactions and amplified with the following thermocycler conditions: an initial denaturation at 95°C for 1 min; 18 cycles of 95°C for 15 sec, 62°C for 15 sec, and 65°C for 2 min; a final extension at 65°C for 10 min; and a 4°C hold. Following amplification, the samples were purified using a 1:1 dilution of AMPure beads as described above.

### Targeted enrichment and preparation of ONT sequencing libraries

Following ligation of the PCR adapters, the four samples had the following concentrations: 36.0 ng/µl (SupCol), 25.2 ng/µl (PFC), 30.2 ng/µl (VCx), and 24.6 ng/µl (caudate) according to genomic DNA screentape analysis (5067-5365 / 5067-5365, Agilent). To prepare the hybridisation reaction, 1 µg of each of the samples was combined with 5 µg Cot-1 DNA (15279011, ThermoFisher) and 1 µl of 1mM blocking oligo (5’ AGGTTAAACACCCAAGCAGACGCCGCAATATCAGCACCAACAGAA 3’) and dried down in a vacuum concentrator for 1h. The dry contents were resuspended by addition of 7.5 µl hybridization buffer and 3 µl hybridization component A (SeqCap EZ Hybridization and Wash Kit: 05 634 261 001, Roche), mixed by tapping, briefly spun by microfuge, and denatured at 95°C for 10 min. The 10.5 µl contents were briefly spun by microfuge and immediately transferred to a pre-warmed 4.5 µl aliquot of the NimbleGen SeqCap EZ Capture probe in a 0.2 ml PCR tube housed in a thermocycler set to 47°C (lid 57°C). Following overnight incubation (∼18 h), 100 µl M-270 Streptavidin Dynabeads (65305, ThermoFisher) per sample were washed twice with 200 µl 1x wash buffer (SeqCap EZ Hybridization and Wash Kit: 05 634 261 001, Roche), once with 100 µl, and then placed in the thermocycler set to 47°C. Incubated samples (15 µl) were transferred immediately to the beads for 45 min. Non-target DNA was removed through washing the beads with buffers of the SeqCap EZ Hybridization and Wash Kit (05 634 261 001, Roche) as per manufacturers instructions with the following modifications: pipette-mixing was replaced by inversion-mixing and brief centrifugation. Beads were finally resuspended in 48 µl nuclease-free water. The following PCR reaction was set up: 48 µl sample; 2 µl PRM primers; 50 µl LongAmp Taq 2x master mix (M0287S, NEB). The reaction mixes were split into 50 µl aliquots and amplified with the following thermocycler conditions: initial denaturation at 95°C for 3 min; 22 cycles of 95°C for 15 sec, 62°C for 15 sec, and 65°C for 10 min; a final extension at 65°C for 10 min; and holding indefinitely at 4°C. Resulting DNA was purified by a 10 min incubation with 70 µl (0.7X) Agencourt AMPure XP beads (A63880, Beckman Coulter), 2x 200 µl washes with 80% ethanol for 30 sec each, 3 min air drying, and elution in 52 µl nuclease-free water. Sample concentrations were determined by Qubit as: 43.5 ng/µl (SupCol), 48.6 ng/µl (PFC), 30.5 ng/µl (VCx), 38.2 ng/µl (caudate).

The captured samples were barcoded using ONT 1D native barcoding genomic DNA kit (EXP-NBD103), and the library prep was performed using the 1D genomic DNA by ligation kit and protocol for the PromethION sequencer (SQK-LSK109). The preparation was performed according to manufacturer recommendations with some modifications. Briefly, each sample underwent end-prep in a 60 µl reaction containing the product from capture, 7 µl Ultra II End-prep reaction buffer, and 3 µl Ultra end-prep enzyme mix. The resulting DNA samples were purified with AMPure beads at a ratio of 1:1 as above. After end-prep, 120 ng of each sample was barcoded by ligation of a barcoding adaptor in a 50 µl reaction containing 2.5 µl of the respective barcode and 25 µl NEB Blunt/TA ligase master mix (M0367). The reaction was incubated at room temperature for 10 min and the resulting product was purified with AMPure beads at a 1:1 dilution as above. Then, 100 ng of each sample was combined into a final reaction to ligate on the sequencing adaptor in a 100 µl reaction containing 400 ng total combined DNA, 25 µl ligation buffer, 10 µl NEBNext Quick T4 DNA ligase, and 5 µl adapter mix. The reaction was incubated for 10 min at room temperature and then purified with AMPure beads at a .55 ratio, to preserve the smaller cDNA fragments, at room temperature for 5 min. The DNA bound to the beads was then washed with and resuspended in 250 µl short fragment buffer and rebound to the magnets twice before being eluted from the beads for 10 min in 25 µl of elution buffer. A PRO002 PromethION flow cell was primed and loaded with the resulting 41.04 ng of captured and barcoded cDNA according to the manufacturer’s recommendation. The LSK109 sequencing protocol was executed, and the sequencing ran for 52 hours.

### Read mapping and clustering of ONT reads

ONT reads were base-called using Guppy base-calling software (ONT) (v1.8.5). Quality control of reads was undertaken using ONT ‘fastq deconcatenate’ (v0.6.2) and POREquality (https://github.com/carsweshau/POREquality). Base-called reads were demultiplexed using Porechop (v0.2.3) (https://github.com/rrwick/Porechop) with the enforced barcode detection parameter. Demultiplexed ONT reads were mapped to a combined index comprising the human genome (hg38) and *in silico* chromosome *(chrIS)*^*^23^*^ using minimap2^36^ (v2.14-r883). The following parameters were used: –ax splice --secondary=no. After retaining only uniquely mapped reads (mapQ=60), reads were processed using the Pinfish suite of tools (https://github.com/nanoporetech/pinfish). First, BAM files were converted into GFF files using the ‘spliced_bam2gff’ tool. We then used the ‘cluster_gff’ tool, which clusters together reads having similar exon/intron structure and creates a rough consensus of the clusters by taking the median of exon boundaries from all transcripts in the cluster. The following parameters were used: –c 3 –d 10 –e 30 –p 1. To assess the performance of ONT in sequencing RNA sequins, we isolated all clustered transcripts aligned to *chrIS* and compared them against the *chrIS* synthetic gene annotations using gffcompare^40^ (v0.10.6), with the following non-default parameters: –M –C –K.

### Illumina short-read sequencing

Samples were spiked with RNA sequins^23^ (Mix A; 2% fractional abundance) before library preparation. RNA CaptureSeq was performed as previously described^28^, using the KAPA Stranded RNA-Seq Library Preparation Kit (Roche). Pre-sequencing quality control qPCR and LabChip GX results were normal. The samples were partitioned into capture pools by RNA yield similarity in order to minimize variation in sample dilution. Sequencing was carried out on an Illumina HiSeq 2500 instrument in high output mode, with 2 × 125 bp paired-end reads.

### Read mapping and assembly of short-read data

Adapters were trimmed from reads using CutAdapt^66^ (v1.8.1) and basic quality control undertaken using FastQC (https://www.bioinformatics.babraham.ac.uk/projects/fastqc/). Reads were aligned using STAR37 (v2.5.3a) to a combined index comprising hg38 and chrIS. STAR was run with the following parameters: --alignMatesGapMax 1000000 --alignIntronMax 1000000 --outFilterIntronMotifs RemoveNoncanonicalUnannotated. Alignments were then assembled into transcript models using Stringtie53 (v 1.3.3), without providing any reference annotations to guide the assembly process. The minimum isoform abundance of predicted transcripts for each locus was required to be at least 1% (–f 0.01). Library type was set as --rf.

### Quantification of gene expression

We compared quantitative gene expression measurements for ONT and Illumina using Salmon^38^ (v0.11.3), providing the *chrIS* annotation and our hybrid transcriptome as a reference. For ONT samples, we used the following non-default parameters: --fldMean 1000 --fldSD 100 --libtype U. For Illumina samples, we used -- libtype ISR. Salmon outputs expression measurements in transcripts per million (TPM).

### Assessment of capture efficiency and quantitative accuracy

Uniquely mapped reads were isolated from BAM files using samtools^67^ (v1.6) ‘view’ (–q 60 for ONT or –q 255 for Illumina). BAM alignments were converted to BED12 files using BEDtools^68^ (v2.25.0) ‘bamtobed’ tool (using the –split flag). We retrieved reads that mapped to targeted regions using BEDtools ‘intersect’; the on-target rate was calculated as the number of reads mapping to targeted regions as a fraction of all reads mapping to hg38.

To assess the performance of CaptureSeq, we plotted the observed abundance (TPM) of each sequin isoform against its input concentration (log_10_ scale). We then carried out simple linear regression of captured and non-captured sequins separately. The limit of detection (LoD) was defined as the spike-in transcript of lowest abundance that was detected in more than one sample. To estimate the enrichment provided by capture, we compared the measured abundance (TPM) of captured vs non-captured standards at each of the shared concentration points. At each shared point, we divided the average TPM for captured standards by the average TPM for non-captured standards, and then averaged all of these to obtain an overall enrichment. Coefficient of variation (CV) was calculated by dividing the standard deviation (SD) of each spike-in by its mean TPM across all four samples. To assess quantitative accuracy at the gene-level, we first counted the number of reads mapping uniquely to annotated sequin loci with featureCounts^69^ (v1.6.3) (using the –L option for ONT data). We then quantified gene expression by calculating the number of reads per gene per 10k reads (RPG10K)^39^ for each sequin locus.

### Assessing the uniformity of sequencing coverage

To assess the uniformity of sequencing coverage, we calculated the read coverage of every base in every sequin exon targeted for capture (*n*=279) using BEDtools^68^ (v2.25.0) ‘coverage’ (with –d and –split options). We then computed the mean and standard deviation (SD) for each exon using BEDtools ‘groupby’, then calculated coefficient of variation (SD / mean) from these metrics. We compared the difference between ONT and Illumina using a paired t-test (2-tailed).

### Generating a hybrid transcriptome

To generate a hybrid transcriptome that leverages both long- and short-read data, we used the FLAIR^41^ tool (v1.2). First, we used FLAIR ‘correct’ by inputting spliced ONT read alignments (from minimap2) and correcting misaligned ONT splice junctions using genome annotations (GENCODE v29; with the –f option) and our accompanying Illumina splice junctions (with the –j option). We used the default window size of 10 bp for correcting splice sites. We then collapsed corrected reads into a non-redundant transcriptome using FLAIR ‘collapse’ with default parameters. This produced a PSL file containing corrected ONT reads, which we converted to BED and GTF files using the UCSC Table Browser https://genome.ucsc.edu/cgi-bin/hgTables).

We then undertook a number of steps in order to produce a high-confidence set of multi-exonic transcripts overlapping GWAS haplotype blocks associated with neuropsychiatric functions. First, we filtered our transcripts using gffread^40^ with the following options: –i 1000000 –U –N –M –K –T. We then annotated our transcripts using gffcompare^40^ (v0.10.6), providing GENCODE v29 as a reference annotation (and using the options –C –K). Any transcript with any the following gffcompare classification codes was removed: ‘e’, ‘p’, ‘r’ or ‘s’. Finally, we removed any transcript that met any of the following criteria: (i) was <200 nt in length, (ii) had <3 exons, or (iii) had no overlap with any capture probe.

### Comparison of our transcriptome to existing annotations

Our set of filtered transcripts was split into unique introns and internal exons using in-house Perl scripts (we disregarded terminal exons because their boundaries are imprecise). We compared these feature counts with the current version of GENCODE (v29), three previous GENCODE releases (v3c, v10 and v19) and MiTranscriptome (v2)^42^. Only multi-exonic transcripts that overlapped targeted haplotype blocks were considered in this analysis. Hg19 annotations were converted to hg38 coordinates using LiftOver^70^.

### Assessing the protein-coding and evolutionary conservation of transcripts

We assessed protein-coding capacity of all assembled transcripts with CPAT^71^ (v1.2.3), using the prebuilt hexamer frequency table and training model (human) obtained from https://sourceforge.net/projects/rna-cpat/files/v1.2.2/prebuilt_model/. For putative coding transcripts (coding probability ≥0.364), open reading frames (ORFs) were predicted and annotated using TransDecoder^72^ (v5.3.0). Protein homology searches were carried out on selected transcripts using the BlastP^73^ and UniProt^74^ databases. Evolutionary conservation of transcripts was investigated using the phastCons 100-way vertebrate alignment^75^; the average score of all nucleotides in each transcript was computed using UCSC’s ‘bigWigAverageOverBed’ script^70^. For both analyses, previously annotated protein-coding transcripts and lncRNAs (GENCODE v29) were included as controls.

### Overlapping transcripts with independent signatures of transcription

We retrieved promoter coordinates from our transcripts using BEDtools ‘flank’ (with –l 1 –r 0 –s options). We downloaded the latest dataset of cap analysis of gene expression (CAGE) robust peaks (hg38) from the FANTOM5 consortium^43^. We used the BEDTools^68^ ‘window’ feature to identify transcripts whose TSS was within 500 bp of a CAGE robust peak on the same strand. We calculated odds ratios of enrichment using the script from Bartonicek *et al*. (2017)^4^. We also overlapped our transcript TSSs with the epigenetic marks that are typically associated with actively transcribed promoters (H3K4me1, H3K4me3 and H3K27ac). We downloaded broad peak calls for these marks (human brain) from the Roadmap Epigenomics Consortium^44^, again lifting them over to hg38 coordinates. Again, previously annotated protein-coding transcripts and lncRNAs (GENCODE v29) were included as controls. Finally, we overlapped our transcripts with a recently published dataset of conserved RNA structures (CRSs) which may be indicative of RNA-mediated functionality^45^.

### Long-range exon coordination analysis

We used our previously developed scripts^33,54^ to look for coordination between distant pairs of exons.

### Visualizing genomic data

Genomic alignments were visualized using the UCSC genome browser^70^ and IGV^76^ (v2.4.10).

### Statistical analyses

Statistical analyses and plotting were carried out using Prism (v7) and R (v3.5.0).

### Data availability

Sequencing libraries have been deposited to GEO with the accession code GSE118158.

## Acknowledgements

We would like to thank Mr James Ferguson and Mr Ted Wong (Garvan Institute) for providing computational support during data analysis. We also thank Dr Ira Deveson (Garvan Institute) for providing access to chromosome 21 CaptureSeq datasets. Brain tissue was received from the New South Wales Brain Tissue Resource Centre at the University of Sydney, which is supported by the Schizophrenia Research Institute. This work was supported by National Health & Medical Research Council (NHMRC) Program Grant 1132524 to GH, JM and CL. SH is supported by a NHMRC Early Career Fellowship (APP1156531). GH is supported by a NHMRC Senior Principal Research Fellowship (1079679). TM is supported by a NHMRC Australia Fellowship (1062470) and also by the Paramor Family Fellowship. HT is a Leon Levy Research Fellow in Neuroscience and is furthermore grateful for a generous gift by Anita Garoppolo. Research reported in this publication was supported by the National Institute on Alcohol Abuse and Alcoholism of the National Institutes of Health under Award Number R28AA012725. The content is solely the responsibility of the authors and does not necessarily represent the official views of the National Institutes of Health.

## Author contributions

JM, SB, GH and CL conceived the study and experimental design. SB, SH, NB and TM designed the RNA capture panel. SH and SB carried out bioinformatics analysis. MS supervised ONT sequencing experiments, bioinformatics analyses and manuscript preparation. JB and KB carried out RNA capture and library preparation for ONT long-read sequencing. DK carried out RNA capture and library preparation for Illumina short-read sequencing. SC processed ONT sequencing data. HT carried out long-range exon coordination analysis. SH and JM wrote the manuscript, with advice from all authors.

