## Supplementary Material for "Targeted, High-resolution RNA Sequencing of Non-coding Genomic Regions Associated with Neuropsychiatric Functions"

**SUPPLEMENTARY MATERIALS**

**Supplementary Figure 1 |** ONT sequencing statistics

**Supplementary Figure 2 |** Read alignment and on-target mapping rates

**Supplementary Figure 3 |** Assessment of quantitative accuracy and capture efficiency (gene-level)

**Supplementary Figure 4 |** Splice junction concordance between ONT and Illumina

**Supplementary Figure 5 |** ONT performance in sequencing full-length synthetic isoforms

**Supplementary Figure 6 |** Improving ONT performance with read clustering

**Supplementary Figure 7 |** Analysis of transcript length and GC content bias

**Supplementary Figure 8 |** Uniformity of sequencing coverage

**Supplementary Figure 9 |** Correlation between ONT and Illumina transcript expression measurements

**Supplementary Figure 10 |** Examples of novel intergenic transcripts validated by CLS study

**Supplementary Figure 11 |** Novel intergenic locus identified on chromosome 7 overlapping GWAS haplotype block associated with multiple sclerosis

**Supplementary Figure 12 |** Novel isoforms of *NRGN* gene overlapping GWAS haplotype block associated with schizophrenia

**Supplementary Figure 13 |** Resolving coordinated splicing events with ONT sequencing

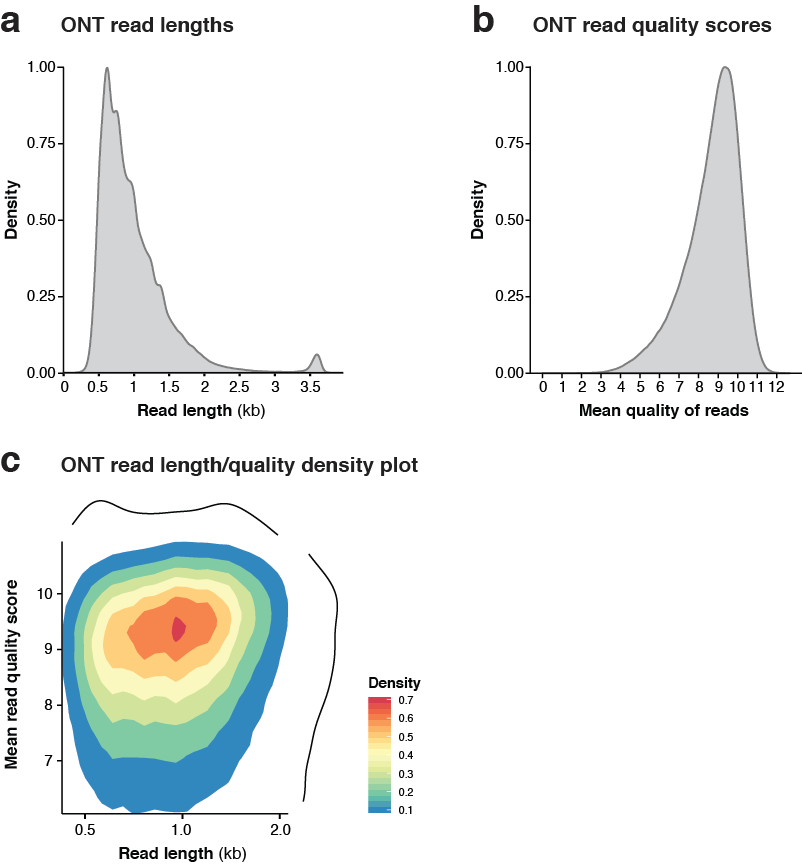

**Supplementary Figure 1 | ONT sequencing statistics**

**(a,b)** Density histograms show distribution of ONT read lengths **(a)** and quality scores **(b)**. **(c)** Density plot shows distribution of ONT reads plotted according to read length (x-axis) and quality score (y-axis). Colours indicate the density of reads.

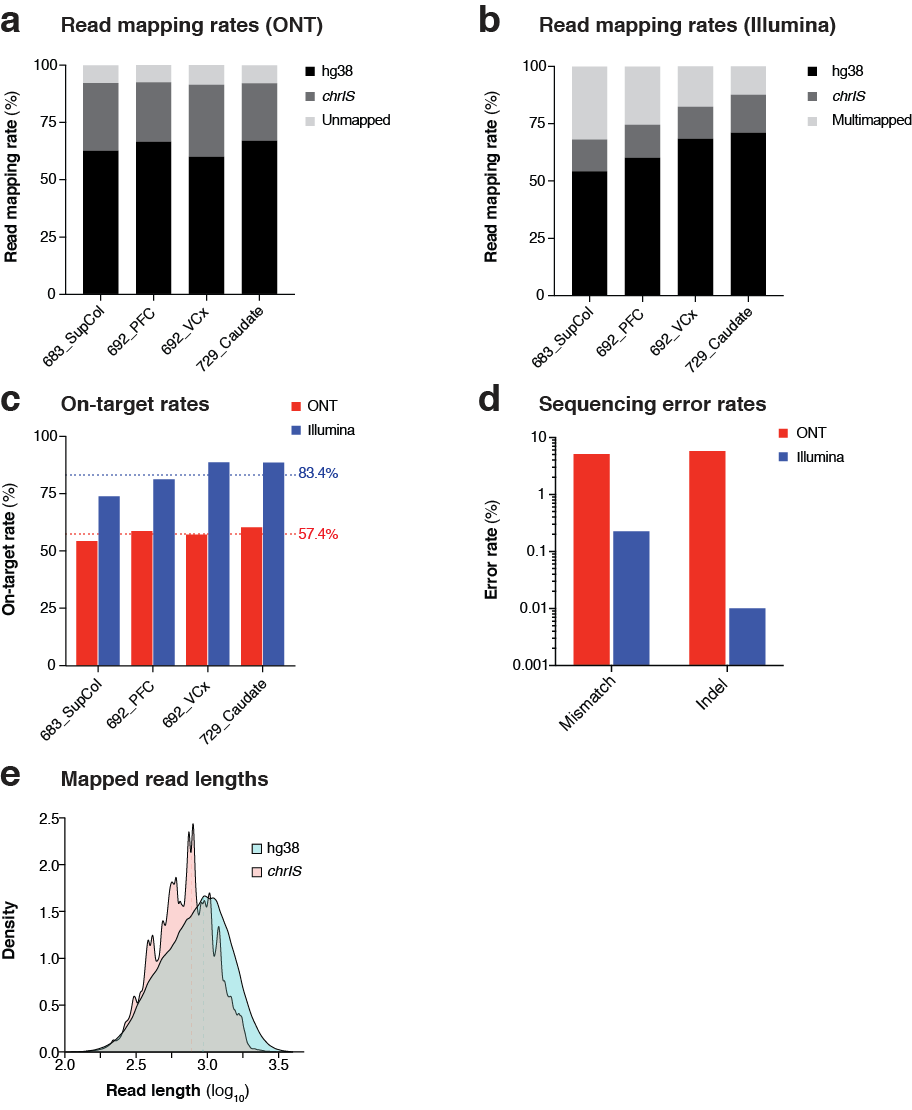

**Supplementary Figure 2 | Read alignment and on-target mapping rates**

**(a,b)** Stacked bar charts show the fraction of reads in each of the four ONT **(a)** and Illumina **(b)** samples which mapped to the human genome (hg38; black), *in silico* chromosome (*chrIS;* dark grey) and those that were unmapped/multimapped (light grey). VCx: primary visual cortex; PFC prefrontal cortex; SupCol: superior colliculus. **(c)** Bar charts show the fraction of reads in each sample that mapped to regions targeted for capture (as a fraction of all hg38-mapped reads). Only reads with perfect mapping scores were considered in this analysis (mapQ=60 for ONT; mapQ=255 for Illumina). ONT samples are shown in red, Illumina in blue. Horizontal dotted lines indicate the mean on-target rate for each technology. **(d)** Bar charts show the mismatch (left) and indel (right) error rates (log_10_ scale) for ONT (red) and Illumina (blue), as calculated using alignments to *chrIS*. **(e)** Density plots show the distribution of mapped read lengths aligning to hg38 (green) and *chrIS* (red). Vertical dashed lines indicate mean.

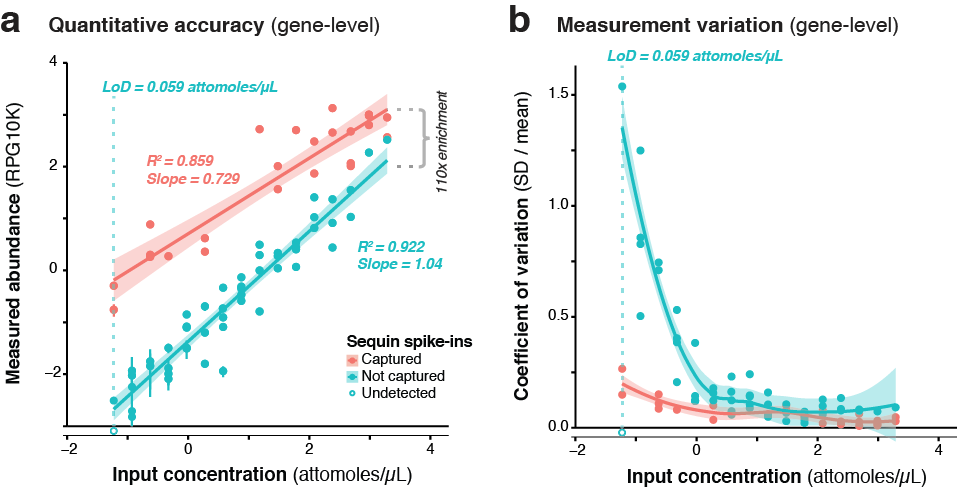

**Supplementary Figure 3 | Assessment of quantitative accuracy and capture efficiency (gene-level)**

**(a)** A subset of RNA sequins were targeted for capture as part of our CaptureSeq design (*n*=25/78 genes). By plotting the measured abundance (reads per gene per 10k reads; y-axis) against the known input concentration (x-axis) for captured (red) and non-captured (green) sequin genes, we can compare the quantitative accuracy of captured vs non-captured transcripts. Open circle indicates the sequin gene that was not detected. Vertical dotted green line indicates the limit of detection (LoD) for non-captured genes. By comparing the difference between the measured abundance of captured and non-captured transcripts, we observe a ~110-fold enrichment of CaptureSeq. Error bars represent standard deviation (SD) between the four replicate ONT samples. **(b)** Scatter plot shows the coefficient of variation (SD divided by mean) of each sequin gene plotted against its respective input concentration, indicating the expression dependent bias of RNA-seq.

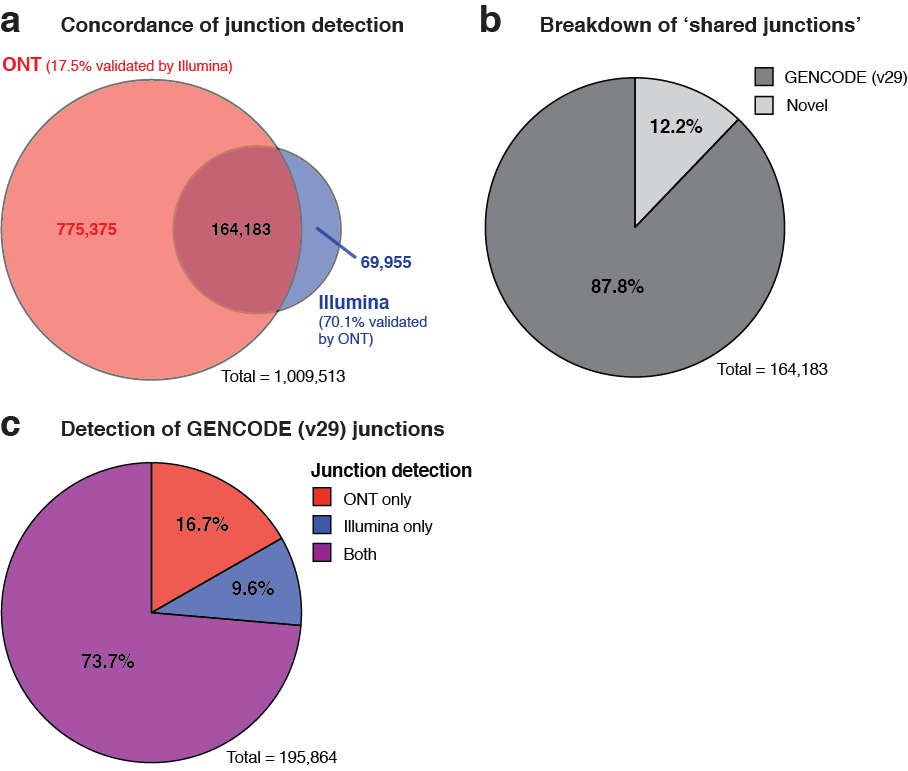

**Supplementary Figure 4 | Splice junction concordance between ONT and Illumina**

**(a)** Venn diagram shows splice junction counts detected by ONT (red) and Illumina (blue). These numbers refer to splice junctions detected with uniquely mapped reads (mapQ=60 for ONT; mapQ=255 for Illumina). **(b)** Pie chart shows how many of the ‘shared junctions’ (i.e. those detected by both sequencing technologies) exactly matched GENCODE (v29) junctions (dark grey) or were novel (light grey). **(c)** Pie chart shows how many GENCODE (v29) junctions were detected by ONT only (red), Illumina only (blue), and by both technologies (purple).

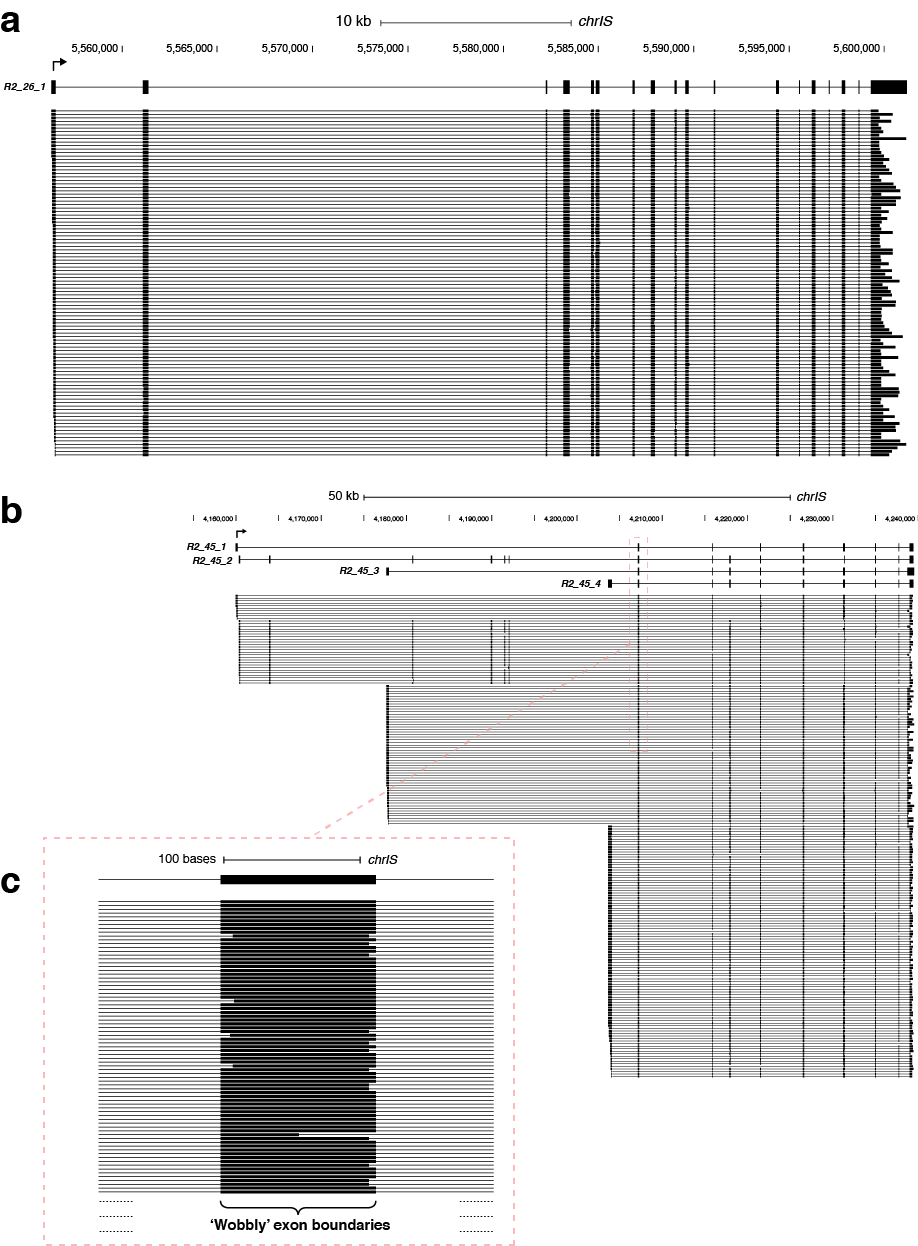

**Supplementary Figure 5 | ONT performance in sequencing full-length synthetic isoforms**

**(a)** Genome browser view showing full-length reads aligned to the *in silico* chromosome (*chrIS*) reference. *R2_26_1* was the longest sequin isoform to be fully sequenced ‘in one go’ (~4.4 kb total length). The top 100 longest full-length reads aligned to this locus are shown. **(b)** Genome browser view showing full-length reads aligned to the *R2_45* synthetic gene locus. ONT sequencing enables the four alternatively spliced isoforms to be unambiguously deconvoluted. **(c)** Magnified view shows a constitutive exon of this locus, with ‘wobbly’ boundaries caused by the relatively high error rate of ONT sequencing. This phenomenon causes high false-positive rates and therefore low precision.

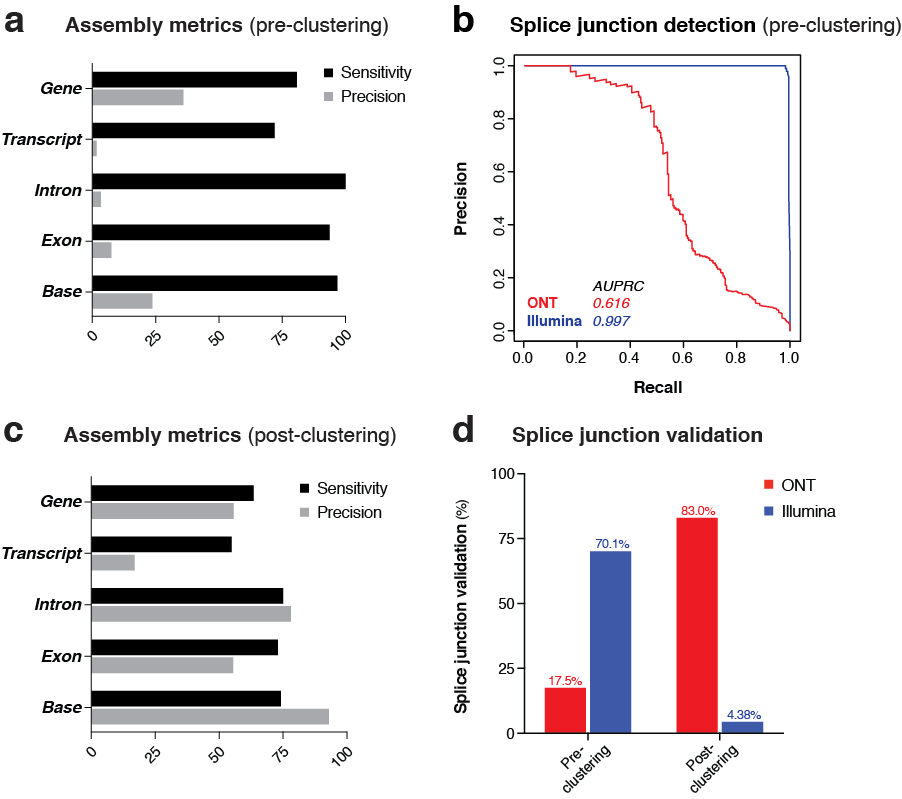

**Supplementary Figure 6 | Improving ONT performance with read clustering**

**(a,c)** Bar charts show the performance of ONT in sequencing RNA sequins in comparison to the *chrIS* annotation, both before **(a)** and after **(c)** clustering. The sensitivity (black) and precision (grey) of assembly are shown at the levels of gene, transcript, intron, exon and base. **(b)** Precision-recall curves compare the performance of ONT (red) and Illumina (blue) sequencing in distinguishing between true-positive (i.e. annotated) and false-positive (i.e. unannotated) splice junctions on *chrIS.* **(d)** Bar charts show the fraction of ONT junctions validated by Illumina (red) and the fraction of Illumina junctions validated by ONT (blue) both before (left) and after (right) clustering.

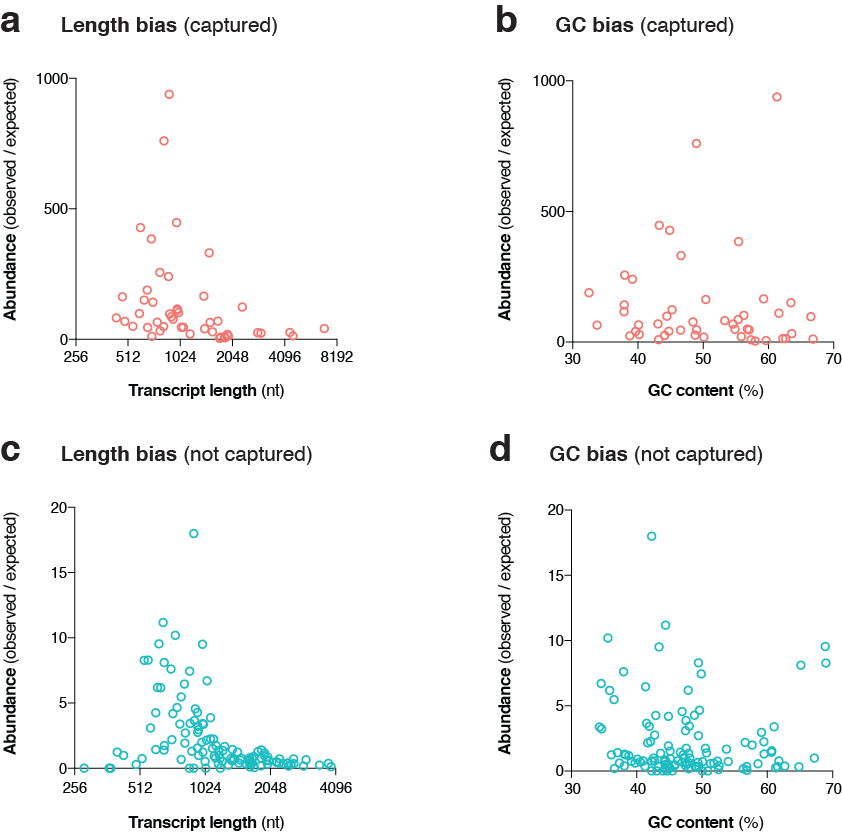

**Supplementary Figure 7 | Analysis of transcript length and GC content bias**

**(a,c)** Scatter plots show the abundance (observed divided by expected) of captured **(a)** and non-captured **(c)** sequin isoforms, plotted against their respective length. **(b,d)** Scatter plots show the abundance (observed divided by expected) of captured **(b)** and non-captured **(d)** sequin isoforms, plotted against their respective GC content.

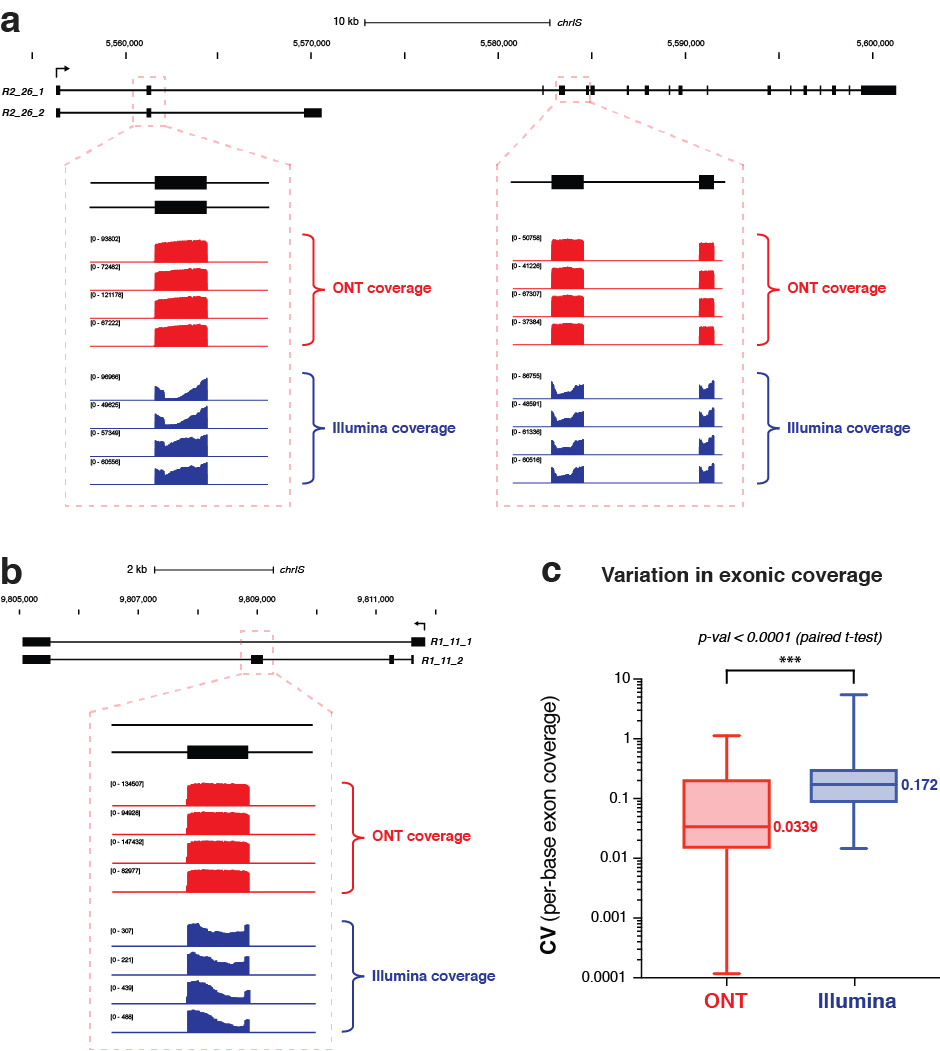

**Supplementary Figure 8 | Uniformity of sequencing coverage**

**(a)** Genome browser view shows synthetic gene locus *R2_26*, with magnified views of selected exons. ONT sequencing coverage (red) of exons was observed to be more uniform than Illumina (blue), as these extreme examples show. Four replicates of each sequencing platform are shown. **(b)** Genome browser view shows synthetic gene locus *R1_11*, with a magnified view of selected exon. ONT sequencing coverage (red) of exon was observed to be more uniform than Illumina (blue). Four replicates of each sequencing platform are shown. **(c)** Box plots show the coefficient of variation (CV; standard deviation divided by mean) of per-base read coverage for each sequin exon targeted for capture (*n=*279 exons). ONT is shown in red, Illumina in blue.

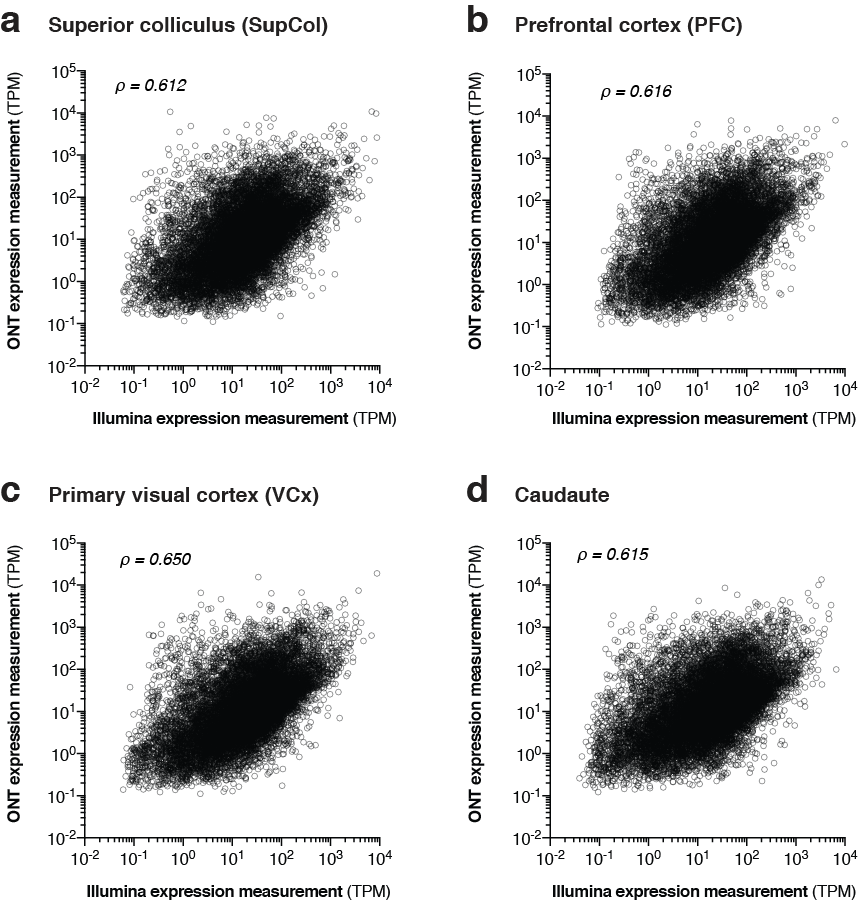

**Supplementary Figure 9 | Correlation between ONT and Illumina transcript expression measurements**

**(a-d)** Scatter plots show the correlation between ONT (y-axis) and Illumina (x-axis) transcript expression measurements (transcripts per million; TPM) using Salmon with our hybrid transcriptome as a reference. Panels show superior collicus **(a)**, prefontal cortex **(b)**, primary visual cortex **(c)**, and caudate **(d)**. Spearman correlation coefficient (ρ) is shown for each sample.

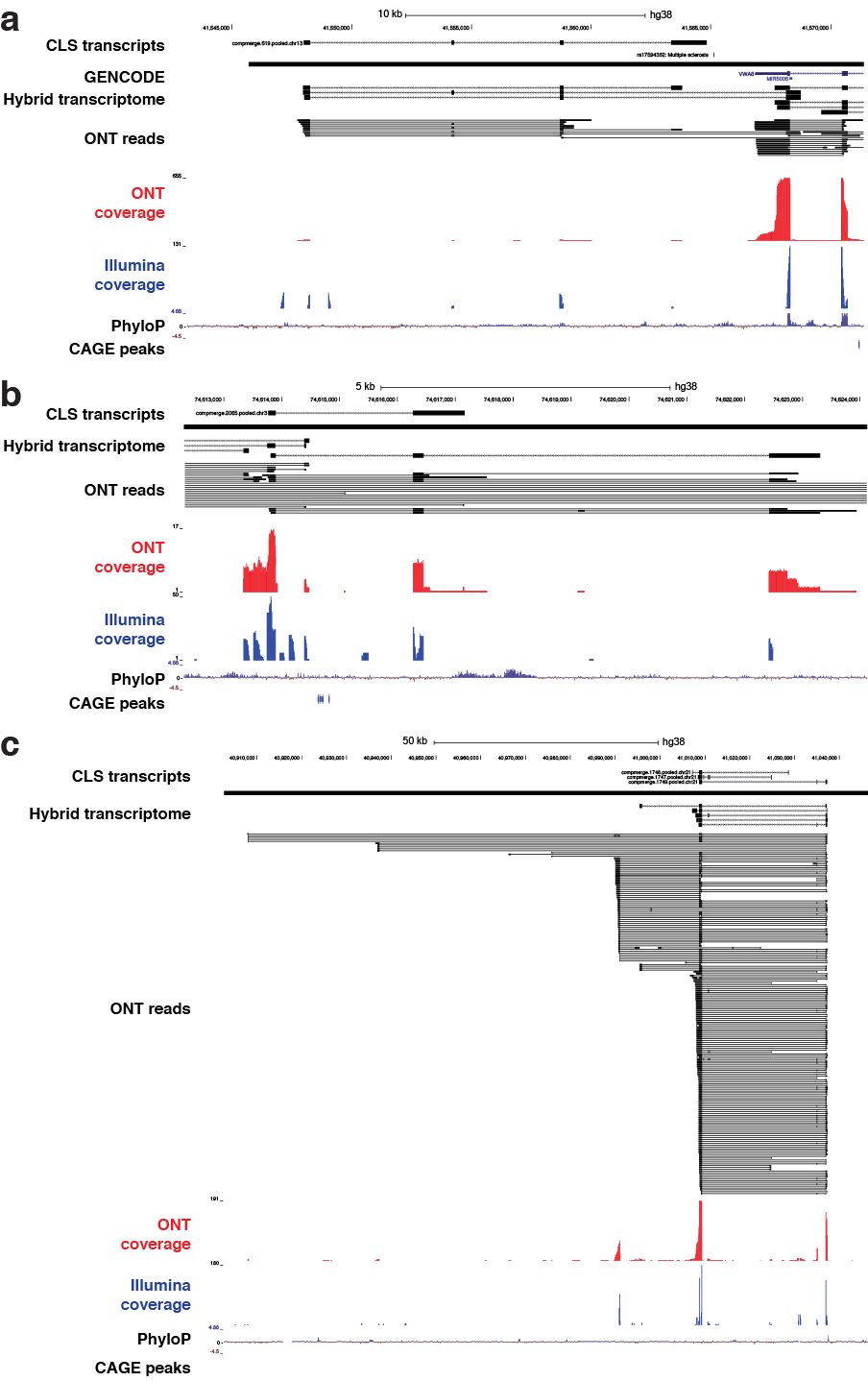

**Supplementary Figure 10 | Examples of novel intergenic transcripts validated by CLS study**

**(a-c)** Genome browser views show examples of our novel intergenic transcripts that, despite not being supported by CAGE peaks at their 5’ ends, nonetheless were independently validated by a recent study that coupled CaptureSeq with PacBio long-read sequencing (CLS). Transcript IDs in ‘CLS transcripts’ track refer to the IDs from the original publication. In each of these examples, our transcriptome expanded upon the PacBio assembly by identifying additional exons and splicing events.

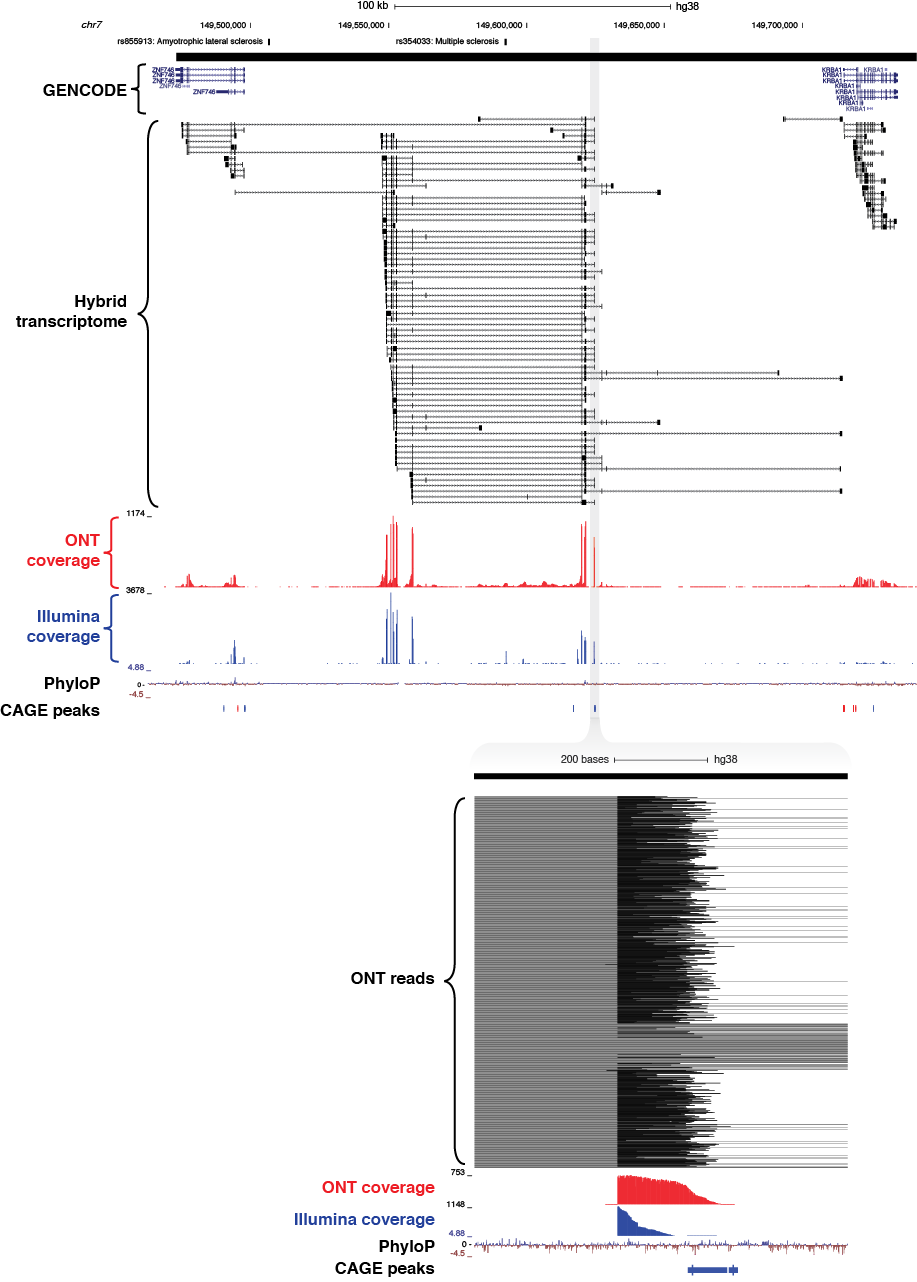

**Supplementary Figure 11 | Novel intergenic locus identified on chromosome 7 overlapping GWAS haplotype block associated with multiple sclerosis**

**(a)** Genome browser view shows a novel intergenic locus identified on chromosome 7 overlapping a ~500 kb GWAS haplotype block (solid black bar, top) associated with multiple sclerosis (rs354033). GENCODE (v29) annotation shown below, followed by spliced ONT sequencing coverage (red), spliced Illumina sequencing coverage (blue), PhyloP conservation track, and CAGE robust peaks. **(b)** Magnified view shows CAGE peak overlapping the novel TSS.

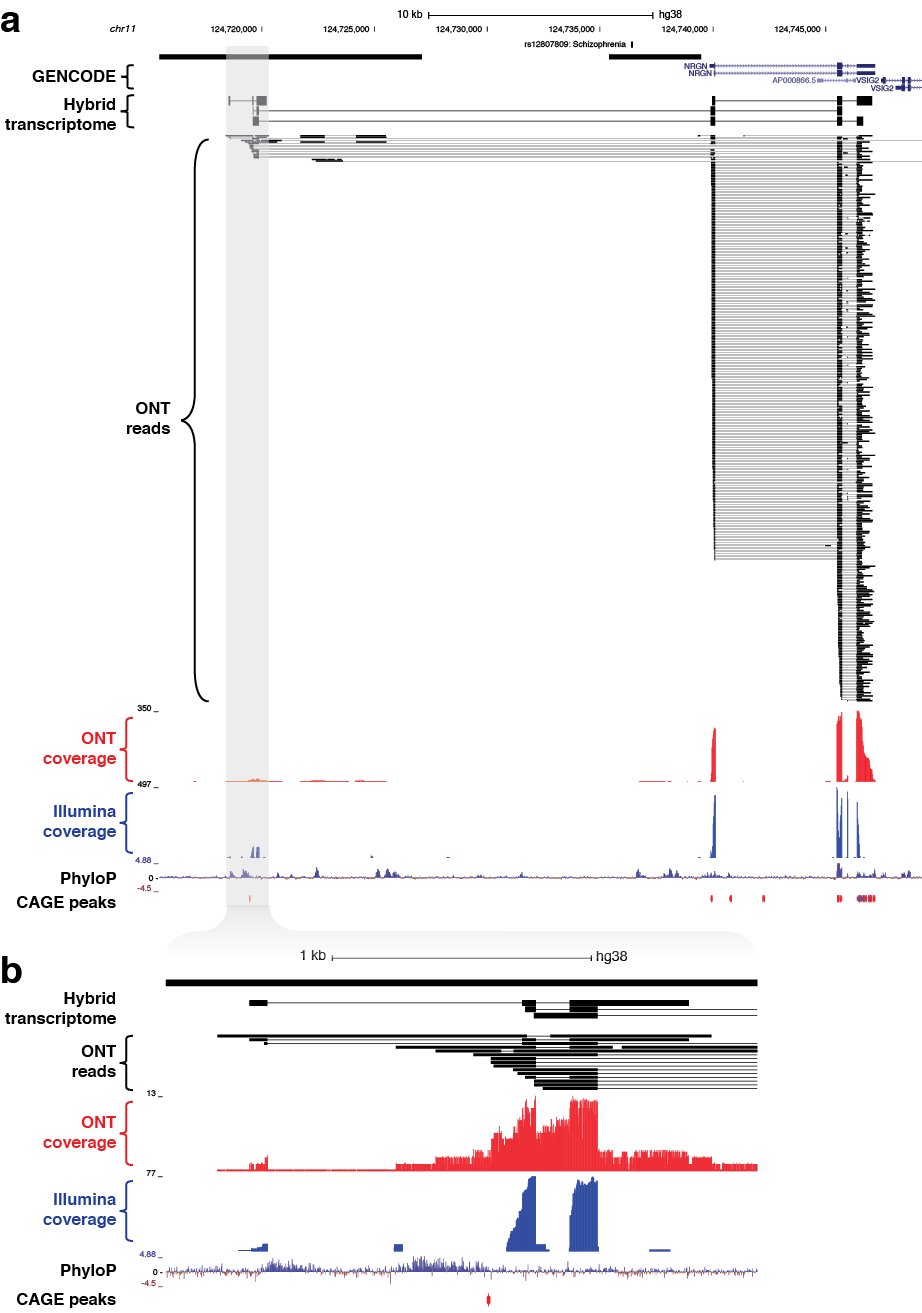

**Supplementary Figure 12 | Novel isoforms of *NRGN* gene overlapping GWAS haplotype block associated with schizophrenia**

**(a)** Genome browser view shows novel isoforms of the *NRGN* gene overlapping a ~4 kb GWAS haplotype block associated with schizophrenia (rs12807809) on chromosome 11. *NRGN* encodes a postsynaptic protein that is thought to be a direct target for thyroid hormone in human brain. **(b)** Magnified view shows the presence of a CAGE peak overlapping the highly conserved promoter region of several novel isoforms.

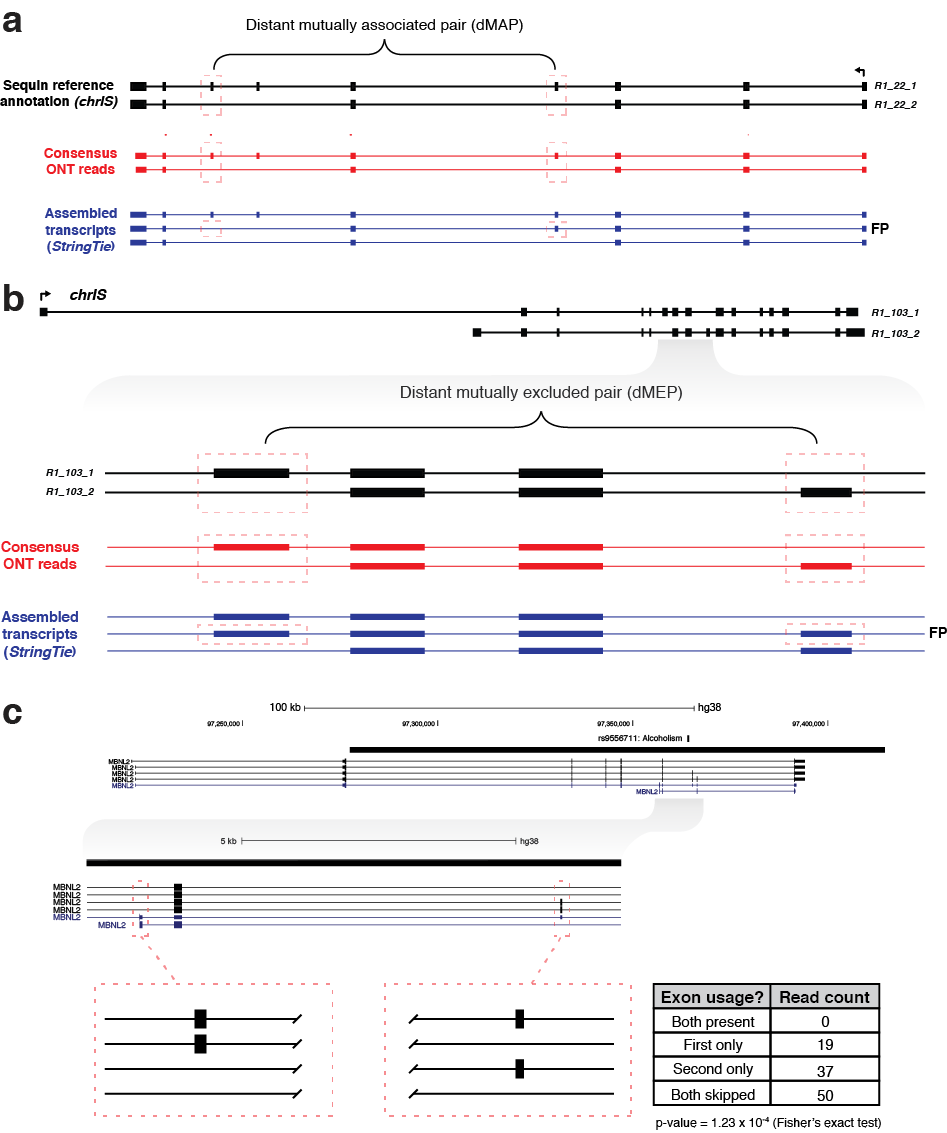

**Supplementary Figure 13 | Resolving coordinated splicing events with ONT sequencing**

**(a)** Genome browser view shows *R1_22* synthetic locus on *chrIS*, which contains a distant mutually associated pair (dMAP) of exons (red boxes). Clustered ONT reads (red) reveal the coordination between these exons unambiguously, with this exon pair always either both present or both absent on a given transcript. However, computational assembly of the Illumina short-read data (blue) produced a false-positive (FP) transcript in which only one of the paired exons is present. **(b)** Genome browser view shows *R1_103* synthetic locus, which contains a distant mutually excluded pair (dMEP) of exons (red boxes). The clustered ONT reads (red) correctly showed that these alternative exons were never present on the same transcript, while the Illumina short-read assembly (blue) produced a FP transcript with both exons present. **(c)** Genome browser view shows coordinated exon pairing in *MBNL2* gene, which overlaps a GWAS haplotype block associated with alcoholism (rs9556711). Two exons in this gene (red boxes) were found to be a dMEP, as they were never both present on the same molecule. Table shows the detected read counts for the four possible isoform scenarios involving these two exons.
